# Intersecting experimental evolution and CRISPR screens to identify novel toxin resistance loci

**DOI:** 10.1101/2025.07.23.666417

**Authors:** Michele Marconcini, Steeve Cruchet, Srishti Goswami, Raghuvir Viswanatha, Matthew Butnaru, Joydeep De, Camilla Roselli, Dafni Hadjieconomou, Norbert Perrimon, Stephanie E. Mohr, Richard Benton

**Author notes:** Corresponding author: E.

## Abstract

Understanding toxin resistance in insects is key to appreciate niche adaptations but remains challenging due to its often-polygenic basis. A well-known example is the specialized association of *Drosophila sechellia* with noni fruit (*Morinda citrifolia*), which is toxic to most other insects, including the closely-related *Drosophila simulans* and *Drosophila melanogaster*. Toxicity of noni is due to its high concentration of octanoic acid (OA), but the mechanisms that determine sensitivity or resistance to OA in different species remain poorly understood. Here, we experimentally-evolved *D. simulans* with increased OA resistance, identifying multiple loci under selection. Cross-referencing these with a genome-wide, OA-resistance CRISPR screen in a *D. melanogaster* cell line highlighted two proteins: Kraken, a putative detoxification enzyme expressed in digestive and renal tissues, and Alkbh7, a mitochondrial protein linked to fatty acid metabolism. Both genes show elevated expression in *D. sechellia* and OA-resistant *D. simulans*. In *D. melanogaster*, *kraken* mutants are more OA-sensitive, while *Alkbh7* overexpression increased OA resistance. Importantly, mutation of these genes in *D. sechellia* reduced OA tolerance. Our identification of genes underlying OA resistance in laboratory and natural contexts demonstrates how complementary, cross-species selection approaches can provide insights into complex mechanisms of toxin susceptibility and adaptation; such methods could also have practical applications in the characterization of natural and artificial insecticides.

## Introduction

How insects resist toxic chemicals in the environment is of both basic scientific interest – reflecting insects’ arms race with plant food sources (Erb and Reymond 2019; Jones, et al. 2022) or venomous predators (Smith, et al. 2013) – and applied interest, because, throughout history, humans have used chemicals to combat insect disease vectors and agricultural pests (Umetsu and Shirai 2020). The best-understood insecticides are those that bind to and modulate neuronal ion channels, where resistance arises through changes in these target proteins (Sparks, et al. 2020; Raisch and Raunser 2023). For example, dichlorodiphenyltrichloroethane (DDT) and pyrethroids prevent closure of voltage-gated sodium channels, resulting in neuronal excitotoxicity and organismal paralysis; diverse insect species displaying resistance for these chemicals have independently-arising mutations in genes encoding these channels (Busvine 1951; Dong 2007). Similarly, neonicotinoids bind to and hyperactivate nicotinic acetylcholine receptors, leading to neuronal death; mutations in various receptor subunits can confer resistance (Matsuda, et al. 2020). Even in such well-studied examples, however, it is thought that other neuronal targets exist (Soderlund and Bloomquist 1989) and/or that the chemicals impact receptors in non-neuronal tissues (Di Prisco, et al. 2013). For many insecticides – as well as other environmental pollutants (Gandara, et al. 2024) – the mode of action is incompletely, or not at all, characterized (Sparks, et al. 2020).

Resistance observed in nature is only partially understood, reflecting the contribution of many genes beyond those encoding target proteins, which can underlie distinct defence mechanisms (Clarkson, et al. 2018; Catteruccia 2020; Lucas, et al. 2023). For example, thickening of the cuticle to reduce entry of insecticides into the body is widely considered an effective resistance mechanism (Balabanidou, et al. 2018; Balabanidou, et al. 2019). Cuticular changes have been linked to overexpression of proteins implicated in exoskeleton development (e.g., cuticle proteins, ABC transporters) but their precise roles, and how they are upregulated, remain unclear (Balabanidou, et al. 2018). Increased expression of metabolic enzymes that degrade insecticides is another commonly observed adaptation in resistant strains, sometimes resulting from copy-number increases in the corresponding genes (Weetman, et al. 2018). Duplicated metabolic genes can also undergo neofunctionalization to produce enzymes with altered specificity, as illustrated by Cytochrome P450s in strains of the brown planthopper that recently evolved resistance to the neonicotinoid imidacloprid (Zimmer, et al. 2018), as well as by more ancient evolution of glutathione S-transferases during transition of the *Scaptomyza* genus of Drosophilidae to herbivory to counter toxic isothiocyanates (Gloss, et al. 2014). However, the lack of experimental resources in many insect species renders it difficult to characterize the causal contribution of specific genetic changes.

A compelling natural model clade to study the evolution of toxin resistance is the trio of closely-related drosophilid species, *Drosophila melanogaster*, *Drosophila simulans* and *Drosophila sechellia* (Fig. 1A). *D. sechellia* is an extreme specialist on the toxic noni fruit of the *Morinda citrifolia* shrub (Jones 2005; Auer, et al. 2021). Amongst *D. sechellia*’s numerous adaptations (R’Kha, et al. 1991; Dekker, et al. 2006; Auer, et al. 2020; Auer, et al. 2021; Alvarez-Ocana, et al. 2023; Shahandeh, et al. 2024), this species has evolved high resistance to octanoic acid (OA), an abundant noni chemical that is toxic to other drosophilids and more divergent insects (Legal, et al. 1994; Jones 1998; Legal, et al. 1999; Markow 2019; Auer, et al. 2021; Kaczmarek, et al. 2022; Kaczmarek, et al. 2024). Previous pioneering studies of *D. sechellia*’s resistance to OA has been studied through genomic mapping-based approaches in *D. sechellia*/*D. simulans* hybrids, which revealed this trait to be polygenic, comprising at least five genomic regions (Amlou, et al. 1997; Amlou, et al. 1998; Jones 1998; Huang and Erezyilmaz 2015). One of the main effect loci encompasses a cluster of 18 genes including members of the *Osiris* (*Osi*) family, which are thought to function in cuticle patterning (Ando, et al. 2019; Sun, et al. 2024). RNA interference (RNAi) knockdown of some *Osi* genes in *D. melanogaster* led to reduced tolerance of OA (Andrade Lopez, et al. 2017; Lanno, et al. 2019). However, genetic evidence for the contribution of these genes to OA resistance in *D. sechellia* is lacking.

**Fig. 1.**
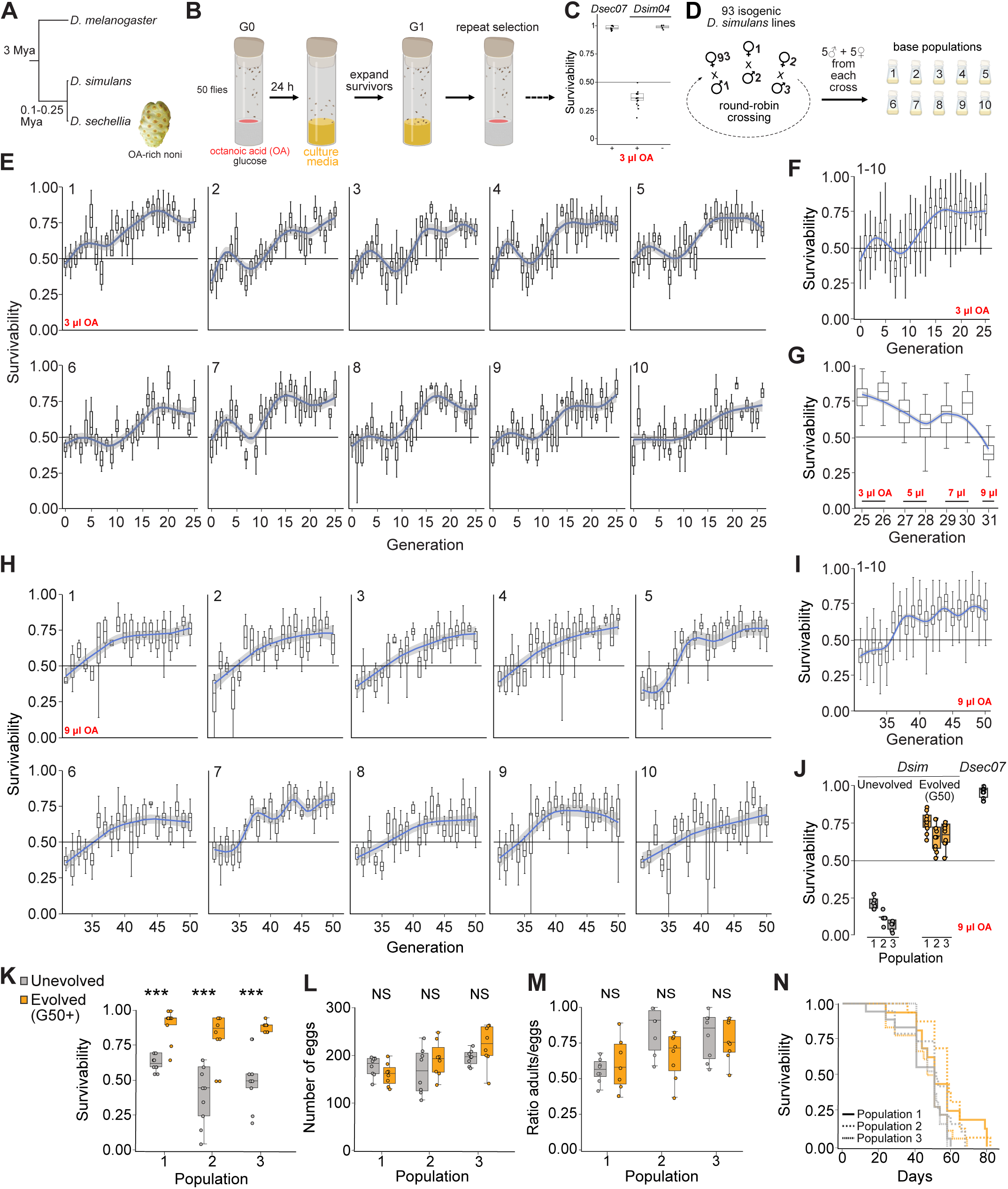
Experimental evolution of *D. simulans* with increased OA resistance. **A** Phylogeny of the drosophilid trio studied in this work. Mya: million years ago. **B** Schematic of octanoic acid (OA) resistance tube assay (see Materials and Methods) and selection procedure. **C** Species differences in resistance of *D. simulans* 04 and *D. sechellia* 07 strains to 3 μl OA. **D** Schematic illustrating the generation of the base populations for Evolve-and-Resequence (see Materials and Methods). **E** Changes in survivability to 3 μl OA of the ten *D. simulans* populations over generations 0-25. Box plots represent the interquartile ranges of the survivability for all the replicates; blue lines shows the generalized additive model (GAM) regression; grey shading represents the 95% confidence interval for the mean responses at each time point. **F** As in **E** for the 10 populations combined. **G** OA resistance for the combined 10 populations with increasing OA volumes during generation 25-31. **H** As in **E**, with 9 μl OA during generations 31-50. **I** As in **H** for the 10 populations combined. **J** Comparison of OA resistance of unevolved *D. simulans* (i.e., the original 10 outcrossed populations maintained on standard fly food; see Materials and Methods), evolved (generation 50 (G50)) *D. simulans*, and *D. sechellia*. **K-N** Phenotypic comparison of unevolved and evolved (G50+) *D. simulans* populations 1-3 for **K** noni resistance (G54), **L** fecundity (G54), **M** developmental viability (G57), and **N** lifespan (G57). For **K-M**, significance was assessed using the unpaired t-test correcting for multiple testing. NS p>0.05, ***p<0.001. For **L-N**, tests were run on standard media. No significant differences between unevolved and evolved flies were detected using the Cox regression model. See Materials and Methods for details and Data S1 for raw data.

Given the challenges of identifying candidate genes through genomic mapping-based strategies, in this work we took two complementary approaches to investigate the evolution of OA resistance in this drosophilid trio: experimental evolution in *D. sechellia*’s closest relative, *D. simulans*, through an Evolve-and-Resequence strategy (Schlotterer, et al. 2016), and CRISPR/Cas9-based genome-wide genetic screening in cultured *D. melanogaster* cells (Viswanatha, et al. 2018). We show how the intersection of these approaches can identify novel genes contributing to OA susceptibility/resistance in these drosophilids.

## Results

### Experimental evolution of OA resistance in *D. simulans*

A previous study demonstrated that *D. simulans* strains can be experimentally evolved for higher OA resistance, enabling the identification of genomic regions (through microsatellite markers) responding to selection (Colson 2004). This success inspired us to take an Evolve-and-Resequence approach, which uses high-throughput sequencing to analyse the entire genome of populations before and after exposure to specific selective pressures, thereby facilitating the identification of adaptive loci from standing genetic variation in controlled and replicable settings (Turner and Miller 2012; Schlotterer, et al. 2016; Barghi, et al. 2019; Kelly and Hughes 2019; Shahrestani, et al. 2021). For studies of insecticide resistance, it has been predominantly used to examine changes in frequency of alleles of known genes rather than as a means to identify novel loci (Zoh, et al. 2021; Sadia, et al. 2024).

The susceptibility/resistance of drosophilids to OA can be easily assessed by exposing flies to the toxin in a culture tube and counting the survivors after 24 h (Fig. 1B). In line with previous observations (Amlou, et al. 1997; Colson 2004), we found that a dose of 3 μl of pure OA (mixed with 1 ml glucose solution), corresponding to roughly half the amount estimated in 2 g of noni pulp (see Materials and Methods), leads to death of ∼60% of *D. simulans* but has no effect on *D. sechellia* (Fig. 1C). By collecting the *D. simulans* survivors, we reasoned that we could allow these to mate and establish the next generation (Fig. 1B), thus enriching genetic variants conferring higher resistance, similarly to (Colson 2004). Repeating the process over multiple generations and multiple replicates would allow us to track phenotypic changes in resistance and investigate the associated trajectories in allele frequencies.

Standing genetic variation should be present in the evolving populations for natural selection to act upon (Teotonio, et al. 2009; Long, et al. 2015). As the spontaneous mutation rate in drosophilids is too low for artificial selection (Keightley, et al. 2014), we generated ten independent polymorphic base populations of *D. simulans* from 93 isogenic strains (Signor, et al. 2018) by round-robin crossing (Fig. 1D). These lines contain millions of SNPs, but we note that they were sampled from a single wild population in California, therefore reflecting the genetic variation present in that locality. The base populations therefore might not include all variants found across the species’ range, including those in regions where *D. sechellia* co-occurs. To minimize bottleneck effects, each population was kept at a size of ∼1000 flies. At each generation, ∼500 flies from each population were randomly collected and selected for resistance to 3 μl OA. Over the course of 25 generations, all the populations showed increased levels of survivability, suggesting parallel adaptive evolution (Fig. 1E). Given the consistent responses across all populations, we also analysed the phenotypic changes grouping all replicates, finding that the mean survivability significantly increased from ∼43% at generation 0 to ∼78% at generation 25 (Fig. 1F).

As we kept the same amount of OA throughout the 25 generations of selection, the selective pressure lowered as the populations’ survivability increased, evidenced as a larger fraction of flies surviving and reproducing. Viewing the phenotype of populations both individually (Fig. 1E) and collectively (Fig. 1F), it was apparent that the change in phenotype plateaued at around generation 15. For continued evolution, we therefore gradually increased the amount of OA during generations 26-31 (Fig. 1G), reaching a selection pressure comparable to the initial phase of the experiment with 9 μl OA (∼38% mean survivability across all populations) (Fig. 1G). We then performed a second round of selection at this dose, observing an increase in survivability across all populations, reaching ∼70% by generation 50 (Fig. 1H). As the populations adapted to the new condition, a second plateau started around generation 40 (Fig. 1H-I). Over 50 generations we were therefore able to evolve *D. simulans* to have approximately 5-fold higher survivability to 9 μl OA compared to the unevolved populations (Fig. 1J), though still falling short of the survival exhibited by *D. sechellia* (Fig. 1J). These plateaus can be explained by the decreasing selective pressure exerted by fixed OA volumes as an ever-greater proportion of flies adapt and survive the treatment. Determining whether an upper limit exists that would constrain the evolution of a *D. sechellia*-like level of phenotypic resistance would require additional selection rounds using progressively higher OA concentrations over an extended (and unknown) number of generations. OA is only one component of noni fruit, although probably the major toxic chemical (Legal, et al. 1994). We therefore tested three of our evolved populations for resistance to noni fruit pulp, finding that they exhibited much higher survival on this substrate compared to the same original populations maintained under neutral conditions (i.e., no OA selection) (Fig. 1K).

Specialization of *D. sechellia* on toxic noni is associated with a reduced egg-laying capacity (R’Kha, et al. 1997; Markow, et al. 2009) and a shorter (albeit strain-specific) lifespan (Watada, et al. 2020; Abe, et al. 2022; Shahandeh, et al. 2024). Our success in evolving *D. simulans* with a partial *D. sechellia*-like resistance trait begs the question as to whether these selected strains exhibit other phenotypic changes as a trade-off for higher toxin resistance. We tested whether the evolved resistant populations of *D. simulans* exhibit similar phenotypic changes in control conditions. However, none of the three focal populations tested exhibited significant decreases in the number of eggs laid (Fig. 1L), developmental viability (Fig. 1M) or lifespan (Fig. 1N). Thus, at least within the timeframe of our experimental evolution experiment, *D. simulans* did not display obvious phenotypic trade-offs for the gain in OA resistance.

### Dynamic directional selection of allele frequencies

We hypothesized that the observed phenotypic changes in *D. simulans* depended on the directional selection of alleles of consistent loci across the ten populations, while most of the genome evolves under neutrality. Allele frequencies at the selected loci should therefore consistently change across all populations, allowing us to statistically distinguish them from randomly evolving loci (Fig. 2A). We sequenced the genomes of the populations of evolved (generation 25 and generation 50) and unevolved (generation 0) flies, identifying 5,127,840 high-quality variants distributed across all chromosomes. Pairwise identity-by-state analysis indicated that as the number of generations increased, the proportions of shared alleles decreased, both between time points and between pairs of populations (left-to-right and top-to-bottom, respectively, in Fig. 2B).

**Fig. 2.**
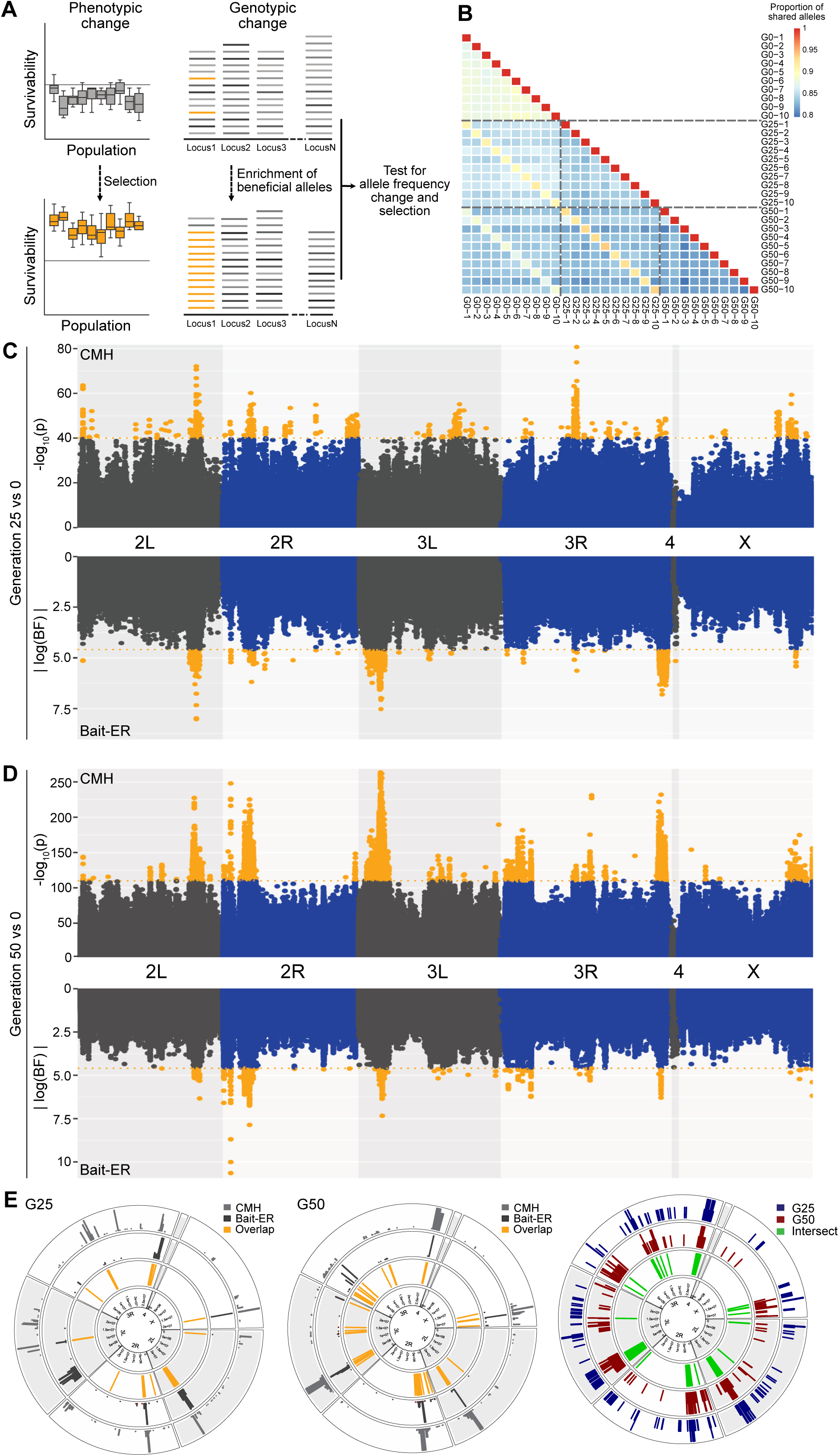
Dynamic directional selection of allele frequencies during experimental evolution. **A** Schematic of the analysis. **B** Identity-by-state analysis. The heatmap depicts the proportion of shared alleles between populations, with values ranging from 0.8 (blue, lowest shared proportion) to 1 (red, identical alleles). **C,D** Manhattan plot showing the probability of allele frequency changes between **C** generations 0 and 25, and **D** generations 0 and 50. Top panel shows the results of the CMH test; orange dots indicate significant variants at a threshold of-log_10_(p) ≥40, a conservative approximation of the top 0.000005% simulated SNPs under neutrality (Fig. S1B). Bottom panel shows the absolute log-Bayes factors estimated by Bait-ER; orange dots indicate significant variants at the conservative threshold log(0.99/0.01). Raw data in Data S2. **E** Circular plots illustrating overlap of predicted genomic regions under directional selection between the two statistical approaches at generation 25 (left) and generation 50 (middle), or between generation 25 and 50 (right). Raw data in Data S3.

To identify the selected genetic variants underlying the phenotypic adaptation, we analysed the genome-wide allele frequency changes both between generations 0 and 25, and between generations 0 and 50, across all populations. The identification of significant changes in frequency is non-trivial, and many methods have been developed (Vlachos, et al. 2019; Kojima, et al. 2020; Barata, et al. 2023). We therefore assessed the significance of frequency changes using two complementary approaches: the Cochran–Mantel–Haenszel (CMH) test (Cochran 1954; Mantel and Haenszel 1959), a widely-used method that tests for allele frequency changes between two time points, and Bait-ER (Barata, et al. 2023), a Bayesian method that tests for selection while explicitly modelling genetic drift. Both tests identified large portions of the genome under positive selection at generation 25 and 50 (Fig. 2C,D). After 25 generations, the two methods shared 1.90 Mb of candidate regions; the overlap encompassed 65.3% of the genomic span identified by CMH and 64.1% of the span identified by Bait-ER. After 50 generations, the two methods shared 4.77 Mb of candidate regions; here, the overlap encompassed 63.6% and 99.9% of the CMH and Bait-ER candidate regions, respectively (Fig. 2E).

Based on the analysis of genome-wide linkage disequilibrium decay (Fig. S1A), we grouped the significant SNPs identified by each of the two methods into genomic blocks according to their proximity (i.e., <50 kb). We retrieved all the candidate genes that fall within the block intervals, obtaining a total of 439 and 401 genes for the CMH and Bait-ER analysis, respectively, at generation 25, and 1238 and 174 genes at generation 50 (Data S3). These genes likely include true targets of selection, but also neighbouring genes affected by hitchhiking. The increase in CMH candidate genes over time – in contrast to the decrease observed with Bait-ER – might reflect the CMH test’s tendency to overestimate significance and to be insensitive to divergence among replicate populations.

The overlap of all significant genes between the two time points is <30%. This might reflect that a large fraction of the candidate genes results from noise caused by genetic drift and draft (hitchhiking) and/or reflect a shift in the phenotypic contribution of genes under the stronger selection (i.e., the higher dose of OA) during generations 27-50. Moreover, previous work on *Drosophila* using temporal sampling showed that adaptation involving complex traits can lead to non-overlapping sets of selected SNPs, in contrast with the simple expectation of continuous increases in allele frequency until fixation (Orozco-terWengel, et al. 2012; Schlotterer, et al. 2016). Gene ontology analysis is complicated at this stage as most hits are likely hitchhikers of the true causal genes. It is also hard to compare currently to previous mapping studies in *D. simulans/D. sechellia* hybrids (Amlou, et al. 1997; Amlou, et al. 1998; Huang and Erezyilmaz 2015) or in the earlier *D. simulans* experimental evolution experiment (Colson 2004), which only identified wide genomic regions. The sole exception, a 170-kb window on 3R containing only 18 genes (including several *Osiris* genes) (Hungate, et al. 2013) did not emerge in our analysis.

### A genome-wide, pooled CRISPR screen for OA susceptibility and resistance genes in *D. melanogaster* S2R+ cells

Given that the previous mapping studies also contain extensive hitchhiking signals and substantial noise across large genomic regions, comparing their results with our analyses would likely yield numerous false positives and still leave a large pool of putative loci, offering limited benefit for homing in on candidate genes. We therefore sought an orthogonal screening approach. Motivated by previous observations of OA toxicity for insect Sf9 cultured cells (Kaczmarek, et al. 2022), we found that OA also leads to reduced viability of *D. melanogaster* S2R+ cells (Fig. 3A). This phenotype prompted us to perform a selection experiment in this cell line through exploitation of a genome-wide, pooled CRISPR/Cas9 mutagenesis approach (Fig. 3B) (Viswanatha, et al. 2018). In brief, a library of sgRNA-encoding plasmids (6-8 sgRNAs/gene) was transfected into Cas9-expressing cells where site-specific plasmid integration predominantly limits targeting to a single gene per cell. Two replicate cultures of this pool of cells were either left untreated or exposed to OA in the medium, at two different concentrations (Fig. 3B). After 26 days in culture, DNA was extracted from the cell pool, then sgRNA sequences were PCR amplified and sequenced to determine those that were enriched after OA selection (reflecting genes whose loss enhances cell viability, referred to hereafter as “susceptibility genes”) or depleted (reflecting genes whose loss leads to lower OA resistance, hereafter “resistance genes”).

**Fig. 3.**
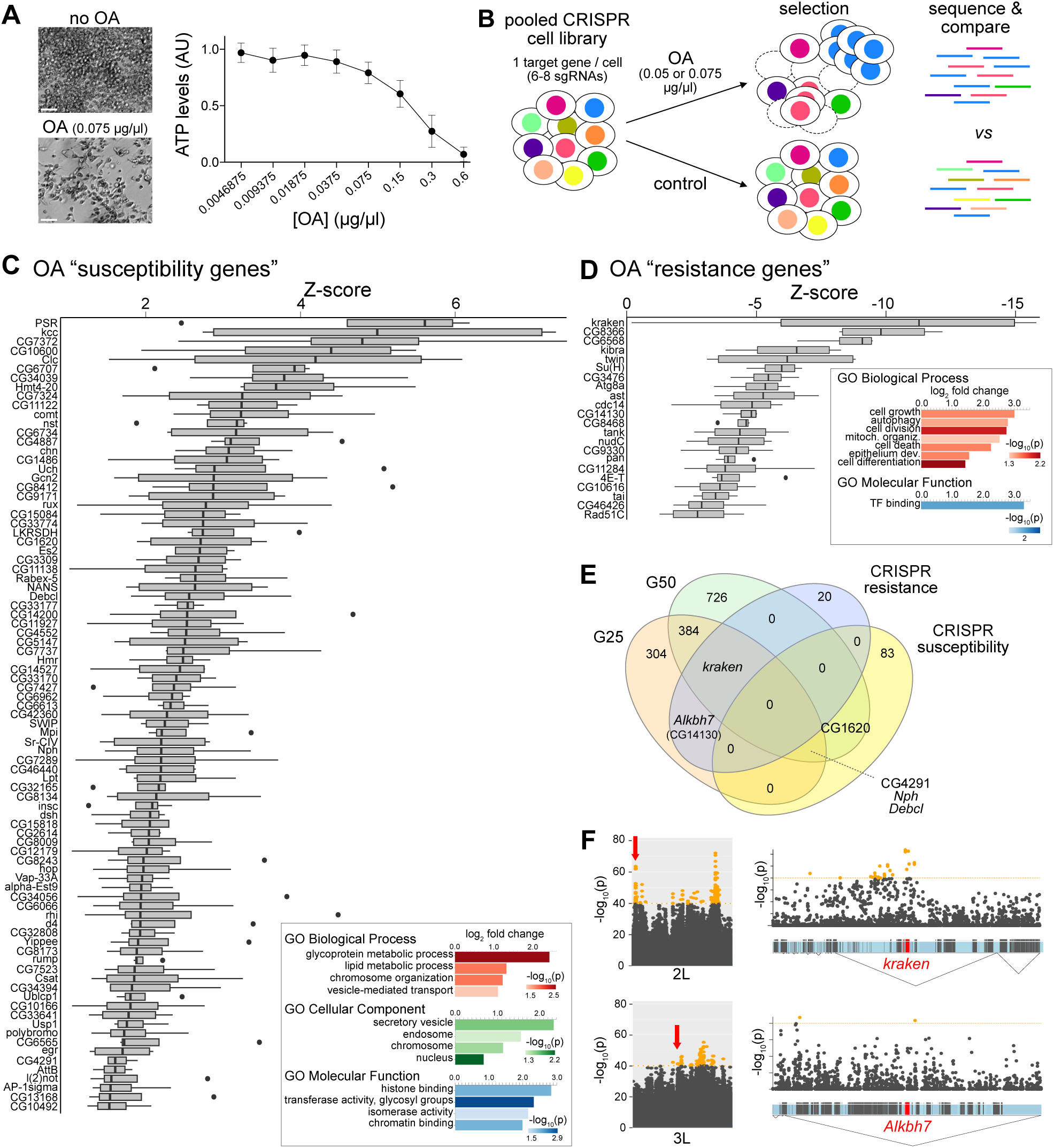
A genome-wide pooled CRISPR screen for OA susceptibility and resistance genes in *D. melanogaster* cultured cells. **A** Left: images of S2R+ cells cultured in the absence or presence of OA. Scale bar, 50 μm. Right: dose-dependent effects of OA on S2R+ cell viability, through measurement of ATP levels with a CellTiter-Glo assay (see Materials and Methods). **B** Schematic of CRISPR screen design. **C** Top 5% “susceptibility” genes (see Materials and Methods and Data S4). Box plots represent the interquartile ranges for all the replicates of the Z-scores obtained through maximum likelihood estimation to test for selection. The inset (bottom-right) displays the gene-ontology analysis of these genes. **D** As in **C**, for the top 5% “resistance” genes. **E** Venn diagram showing the overlap between all the genes identified in the experimental evolution at G25 and G50, and genes associated with susceptibility and resistance in the CRISPR screen. **F** Subportion of the top panel (CMH test) of the Manhattan plot from Fig. 2C for chromosome arms 2L and 3L, with the red arrows highlighting the regions magnified on the right showing the significant variants and linkage disequilibrium blocks (triangles) for *kraken* and *Alkbh7* (CG14130).

To identify candidate OA susceptibility/resistance genes, we first filtered out genes essential for general cell survival (Viswanatha, et al. 2018) and genes that were not detectably expressed in S2R+ cells (using DGET (Hu, et al. 2017)). (Removal of such genes are inevitable limitations of this cell-based screening approach; see Discussion). After comparing results from each condition and replicate, we identified the top 5% susceptibility genes (Fig. 3C) and top 5% resistance genes (Fig. 3D). Amongst the 87 susceptibility genes, there was no single stand-out hit, suggesting that – at least in S2R+ cells – OA does not have a single major cellular target (unlike, for example, insecticidal bacterial toxins that bind a specific cell surface receptor (Xu, et al. 2022)). Gene-ontology analysis of these susceptibility genes revealed a significant enrichment of genes involved in membrane trafficking (Fig. 3C). As OA has previously been hypothesized to disrupt cellular membranes (Borrull, et al. 2015; Kaczmarek, et al. 2022), the recovery of these hits suggests that general reduction in membrane transport reduces OA uptake into cells and/or renders membranes less susceptible to OA treatment. 22 OA resistance genes were identified, encoding diverse types of proteins (Fig. 3D), suggesting that there are multiple mechanisms by which cells can resist the toxic effects of OA.

We next examined which of the genes identified in the S2R+ cell selection experiments might be relevant for evolution of OA resistance in whole animals by determining those also present within significant blocks of the experimental evolution analyses (Fig. 3E,F). Four susceptibility genes were identified: CG1620 (a predicted SANT-MYB domain transcription factor), CG4291 (a putative component of the splicing machinery), *Nph* (a chromatin remodelling factor (Emelyanov, et al. 2014)) and *Debcl* (a pro-apoptotic member of the Bcl-2 family (Galindo, et al. 2009)); these are all likely to have indirect roles in conferring susceptibility to OA given their broad functions. By contrast, two resistance genes have potentially direct roles in conferring tolerance to OA: *kraken* and CG14130. *kraken* encodes a putative serine hydrolase (Edwin Chan, et al. 1998) and stood out both because it was the top hit in the resistance gene screen (Fig. 3D), and because of its location in one of the most prominent peaks in the experimental evolution at generations 25 and 50 (Fig. 3F and Fig. 2C-D). CG14130 encodes the orthologue of mammalian *Alkbh7*, which is involved in regulating fatty acid metabolism (Solberg, et al. 2013; Zhang, et al. 2021), and is located within a significant peak at generation 25 (Fig. 3F and Fig. 2C). Therefore, intersecting the experimental evolution and CRISPR screening results allowed us to narrow down hundreds of potential candidates to two focal genes for downstream investigations.

### Sequence, expression and evolutionary analyses of candidate OA resistance genes

To assess the candidacy of *kraken* and CG14130 (hereafter *Alkbh7*) in contributing to OA resistance in animals, we first examined their sequence and expression in *D. melanogaster*. Kraken belongs to a large family of serine hydrolases, which encompass lipases, esterases and proteases that use a nucleophilic serine residue in the active site for hydrolysis of substrates (Edwin Chan, et al. 1998). Consistent with the conservation of this serine in Kraken (Fig. 4A and Fig. S2A), an activity-based proteomic screen of the serine hydrolase superfamily in *D. melanogaster* provided evidence for *in vivo* catalytic activity of Kraken (Kumar, et al. 2021). *kraken* is broadly expressed in adults, with particular enrichment in the gut and Malpighian tubules (Fig. 4B), similar to previous observations of *kraken* expression in the embryo by RNA *in situ* hybridization (Edwin Chan, et al. 1998). Using the Fly Cell Atlas (Li, et al. 2022), we examined sub-tissue adult expression of *kraken*: in the Malpighian tubules, *kraken* is expressed at highest levels in the principal cells (Fig. 4C), which are involved in ion transport and detoxification. In the gut, *kraken* transcripts are most robustly detected in various types of enterocytes, which secrete digestive enzymes (Lemaitre and Miguel-Aliaga 2013) (Fig. 4C). However, Kraken lacks an N-terminal signal sequence (Teufel, et al. 2022), suggesting that the protein remains in the cytoplasm of these cells, unlike canonical digestive enzymes (Lemaitre and Miguel-Aliaga 2013), and more similar to detoxification enzymes of the Cytochrome P450 and Glutathione S-transferase families (Yang, et al. 2007). To examine *kraken* expression *in vivo*, we performed RNA fluorescence *in situ* hybridization, which detected *kraken* transcripts in various region of the gut (crop, crop duct, proventriculus, hindgut, ampulla) and Malpighian tubules (Fig. 4D).

**Fig. 4.**
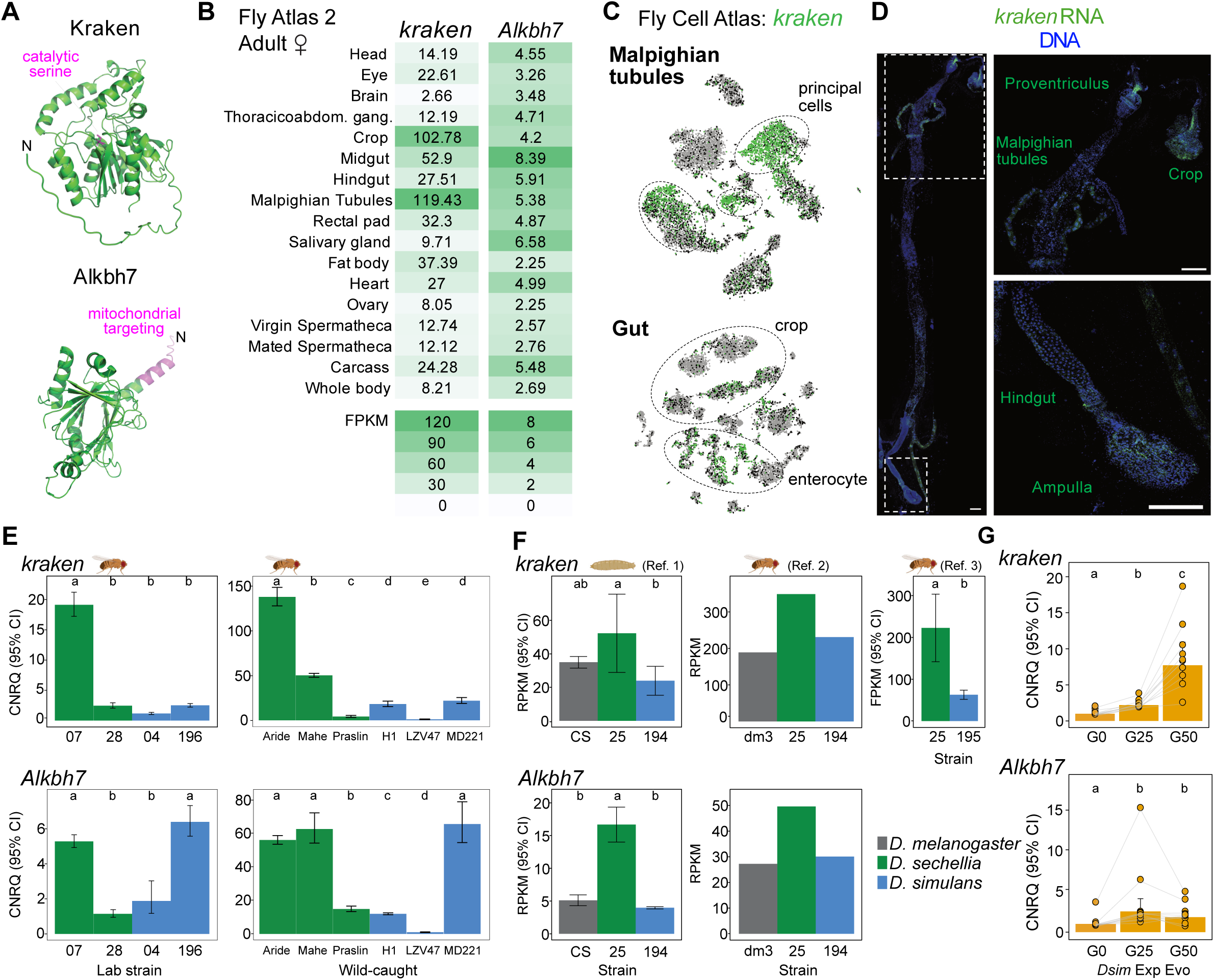
Sequence and expression analyses of *kraken* and *Alkbh7*. **A** AlphaFold protein models for *D. melanogaster* Kraken (AF-O18391-F1-model_v4) and Alkbh7 (AF-Q9VTP1-F1-model_v4) (Jumper, et al. 2021; Varadi, et al. 2022), highlighting a conserved putative catalytic serine in the active site in Kraken, and the mitochondrial targeting sequence in Alkbh7 (predicted by MitoFates (Fukasawa, et al. 2015)). See also Fig. S2A,B. **B** Tissue-specific expression of *D. melanogaster kraken* and *Alkbh7* from bulk RNA-sequencing data from the Fly Atlas 2.0 (Krause, et al. 2022). **C** tSNE plots illustrating *D. melanogaster kraken* expression in single-cell transcriptomes of the Malpighian tubules and gut from the Fly Cell Atlas (Stringent 10x datasets) (Li, et al. 2022). **D** Left: RNAscope detection of *kraken* (green) transcripts in the gut and Malpighian tubules, with nuclei counterstained with DAPI (blue). Right: higher-magnification images showing *kraken* transcript expression in the indicated tissues. Scale bars, 200 μm. This expression pattern was observed in tissues from >20 individuals. **E** Expression levels of the *kraken* and *Alkbh7* in the indicated laboratory strains (left panel) and wild caught strains (right panel) of adult female *D. sechellia* and *D. simulans* measured by qPCR (Data S6). Expression is represented as calibrated and normalized relative quantities (CNRQ). Significance was assessed using the unpaired t-test correcting for multiple testing and represented using letter codes. **F** Expression levels of *kraken* and *Alkbh7* in the indicated strains and species at larval or adult stages as indicated from published RNA-seq data. (References: 1, (Watanabe, et al. 2019), 2 (Ma, et al. 2018), 3 (Kalra, et al. 2024)). Significance was assessed using the unpaired t-test correcting for multiple testing and represented using letter codes when multiple replicates were present in the original data. **G** Expression levels of the candidate genes at generation 0, 25 and 50 of the experimentally-evolved *D. simulans* populations measured by qPCR (Data S6). Significance was assessed using the paired t-test correcting for multiple testing and represented using letter codes.

Alkbh7 is a member of the alpha-ketoglutarate-dependent hydroxylase (Alkbh) family (Fig. 4A), which comprises enzymes that act on diverse substrates in processes such as biosynthesis, metabolism and epigenetic regulation (Xu, et al. 2021). As in mammals (Solberg, et al. 2013), *D. melanogaster Alkbh7* displays broad tissue expression (Fig. 4B). Murine Alkbh7 is localized to the mitochondrial matrix (Solberg, et al. 2013). This property appears to be conserved in the *Drosophila* protein, which has a mitochondrial targeting sequence (Fig. 4A and Fig. S2B) (Fukasawa, et al. 2015); consistently, we identified Alkbh7 within *D. melanogaster* proteomes of the mitochondrial matrix (Chen, et al. 2015; Sen and Cox 2022).

Although the sequences of both Kraken and Alkbh7 are overall highly similar across *D. melanogaster*, *D. simulans* and *D. sechellia* (Fig. S2A,B), we wondered whether we could detect signs of selection at these loci in *D. sechellia* employing population sequence datasets of *D. sechellia* and *D. simulans* (Schrider, et al. 2018). Using the McDonald-Kreitman test (McDonald and Kreitman 1991) on the coding sequences of both genes, we found evidence for gene-level positive selection on *kraken* but not *Alkbh7* (Fig. S2C). Because selection might act pervasively at individual sites rather than along the entire protein coding region, we also implemented a codon-based analysis using FUBAR (Fast, Unconstrained Bayesian AppRoximation) (Murrell, et al. 2013), which identified 1 and 4 codons under diversifying positive selection in *kraken* and *Alkbh7,* respectively (Fig. S2D). Although their functional relevance remains unclear, such mutations might reflect underlying evolutionary dynamics at the protein level.

We next compared the expression levels of these genes both in laboratory and wild-caught strains of *D. sechellia* and *D. simulans* (Fig. 4E), as well as in RNA-seq datasets of various strains of *D. sechellia*, *D. simulans* and *D. melanogaster* (Ma, et al. 2018; Watanabe, et al. 2019; Kalra, et al. 2024) (Fig. 4F, Fig. S3A,B). While there were some exceptions, overall both *kraken* and *Alkbh7* show higher expression levels in *D. sechellia* adults and larvae compared to the other species. Importantly, in our experimentally-evolved *D. simulans* populations, we also found that both *kraken* and *Alkbh7* were more highly expressed compared to unselected populations (Fig. 4G). The expression differences between species, and between unevolved and evolved *D. simulans*, reflect variation in basal levels of expression, as prior exposure to OA did not lead to changes in expression in either *D. sechellia* or *D. simulans* (Fig. S3C). RNA FISH of *kraken* in *D. sechellia* revealed similar spatial expression as in *D. melanogaster*, with the addition of a robust signal in the midgut (Fig. S3D). These observations suggest that elevated expression in this species is due, at least in part, to a novel expression pattern in this digestive organ.

### Functional contributions of *kraken* and *Alkbh7* to OA resistance

Given the expression differences observed in the naturally and experimentally evolved OA-resistant drosophilids, we asked whether artificially modulating gene expression would impact OA tolerance. We first performed ubiquitous RNAi of these genes in *D. melanogaster* (Fig. S4A) and assessed the effects on resistance to OA using a short-term, plate-based assay (Jones 1998) (see Materials and Methods). *kraken^RNAi^* led to significantly diminished OA resistance compared to genetic controls, while *Alkbh7^RNAi^*did not have a discernible effect (Fig. 5A). We next generated deletion mutants for these genes in *D. melanogaster* (Fig. S4B). Similar to the RNAi experiments, *kraken* mutants, but not *Alkbh7* mutants, displayed reduced resistance to OA (Fig. 5B). All genotypes remained fully viable when no OA was added (Fig. S5A).

**Fig. 5.**
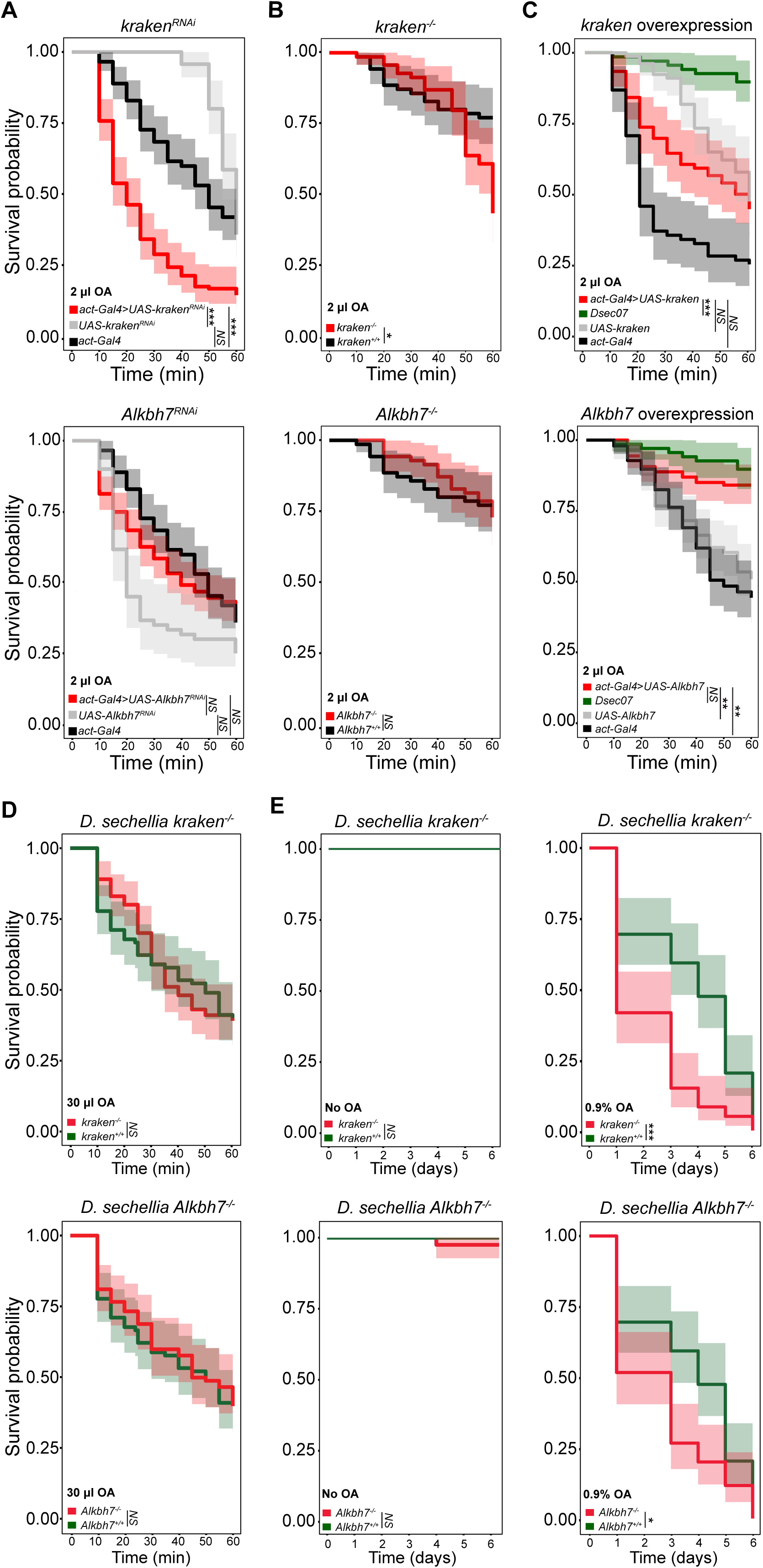
Functional validation of the contribution of *kraken* and *Alkbh7* to OA resistance. A-C. Survival curves of adult flies of the indicated genotypes tested in the plate assay with 2 μl OA for *kraken* and *Alkbh7* RNAi (**A**), mutants (**B**) and overexpression (**C**). Each replicate consisted of 10 2-7-day-old female flies. N replicates ≥6, n flies ≥60. Shading indicates 95% confidence intervals. Genotypes: **A** *act-Gal4/+;UAS-kraken^RNAi^/+, UAS-kraken^RNAi^/+, act-Gal4/+, act-Gal4/+;UAS-Alkbh7^RNAi^/+, UAS-Alkbh7^RNAi^/+, act-Gal4/+* **B** *kraken^1^*, *Alkbh7^1^, Canton-S* (wild-type genetic background control for both mutants), **C** *act-Gal4/+;UAS-kraken^o/e^/+*, *UAS-kraken^o/e^/+*, *act-Gal4/+*, *act-Gal4/+;UAS-Alkbh7^o/e^/+*, *UAS-Alkbh7^o/e^/+*, *act-Gal4/+*. Significance is based on the resulting mixed effect Cox regression model. NS p>0.05, *p<0.05, **p<0.01, ***p<0.001. The same data for *D. sechellia* are shown in both plots in **C**. **D** Survival curves of wild-type (*Dsec07*), and *kraken* (*Dsec\kraken^RFP^*) and *Alkbh7* (*Dsec\Alkbh7^RFP^*) mutant *D. sechellia* in the plate assay with 30 μl OA. Statistics as in **A**. **E** Long-term survivability of the same genotypes as in **D** (n = 60 per condition) in control conditions (no OA) (left) and food supplemented with 0.9% OA (right). Statistics as in **A**. Raw data in Data S7.

Loss-of-function analyses might fail to uncover a role for *Alkbh7* in whole animals if there are redundant resistance mechanisms absent from S2R+ cells. We therefore took a complementary approach by overexpressing these genes in *D. melanogaster*. Strikingly, overexpression of *Alkbh7* resulted in survivability of *D. melanogaster* to levels close to those observed for *D. sechellia* under the same experimental conditions (Fig. 5C, Fig. S4C), indicating that this gene is sufficient to confer high-level OA resistance. By contrast, *kraken* overexpression did not have any detectable impact on OA resistance (Fig. 5C); a similar lack of effect was seen when overexpressing *D. sechellia kraken* (Fig. S2E). The absence of a phenotypic effect upon *kraken* overexpression is surprising, given its elevated expression in *D. sechellia* and in evolved *D. simulans* but this could reflect either insufficient overexpression levels (Fig. S4C) and/or genotype-by-genotype interactions that are not recapitulated when manipulating a single locus in a *D. melanogaster* background.

A key question is whether these genes contribute to OA resistance of *D. sechellia*. Implementing CRISPR/Cas9-based genome-engineering in this species (Auer, et al. 2020; Auer, et al. 2021), we generated and validated null mutants in these genes in *D. sechellia* (Fig. S4B). These mutant flies were homozygous viable and fertile, with no obvious morphological or behavioural abnormalities, allowing us to test their resistance to OA in the plate assay. Although we did not detect any significant difference to genetically-matched wild-type flies (Fig. 5D), this was anticipated given the polygenic nature of OA resistance in *D. sechellia* (Amlou, et al. 1997; Amlou, et al. 1998; Jones 1998; Huang and Erezyilmaz 2015), and the modest or absent phenotypes of equivalent mutants in *D. melanogaster* (Fig. 5B). Likewise, rearing *D. sechellia* null mutants on noni pulp (Fig. S5B), we observed no significant differences between mutant and wild-type flies over >30 days, indicating that loss of either gene alone does not compromise *D. sechellia*’s capacity to persist in its natural host fruit.

We reasoned that more subtle differences might be detected by monitoring *D. sechellia* survivability at higher OA concentrations over several days. We therefore exposed flies to food containing 0.9% OA, approximately threefold higher than the ∼0.3% OA reported for ripe noni fruit (3.06 g OA/kg fruit) (Pino, et al. 2009). Indeed, we observed higher mortality rates of both *kraken* and *Alkbh7* mutants compared to genetically matched wild-type flies over a period of 6 days (Fig. 5E). By contrast, both mutants survived equally well as controls in the absence of OA over this time period (Fig. 5E). To determine whether the observed OA sensitivity was attributable to a general reduction in fitness, we examined survivorship under several conditions of thermal stress (Fig. S5C): mutant and wild-type flies showed no significant differences across all temperatures tested, indicating that the increased mortality observed upon OA exposure does not result from an overall physiological weakness of the knockout lines. These data provide the first evidence for the contribution of specific genes to OA resistance in *D. sechellia*. However, future reciprocal hemizygosity tests (Stern 2014) and/or allelic swap studies with *D. simulans* or *D. melanogaster* would be required to assess if and how they contribute to species-specific differences in OA sensitivity (e.g., through structural and/or regulatory differences in the genes).

## Discussion

Identifying specific loci underlying phenotypic evolution is challenging due to the polygenicity of the vast majority of adaptive traits, with recent advocacy of new experimental paradigms to bridge genotype-phenotype relationships (Tautz, et al. 2026). Investigating the evolution of chemical toxicity is an attractive model trait because of the relatively easy phenotypic read-out. To gain insight into the genetic basis of susceptibility and resistance to a natural toxin, OA, we have intersected the results of selection over vastly different timescales: weeks in cell lines, years in experimentally-evolved laboratory strains, and millennia during speciation.

Unlike insecticides that activate or antagonize broadly-expressed neuronal ion channels, the mode of action of OA appears to be mechanistically very different. Beyond insect cell lines (our work and (Kaczmarek, et al. 2022)), OA is also toxic to budding yeast (Mota, et al. 2024) and bacteria (Liu, et al. 2013), suggesting that this toxin acts via a more fundamental cellular mechanism. Indeed, a genome-wide, knock-out screen in yeast for genes required for tolerance of OA highlighted contributions of several core cellular processes, such as vesicle trafficking, mitochondrial function or chromatin regulation (Mota, et al. 2024), reminiscent of our CRISPR screen results. Consistent with this idea, the results of our experimental evolution and CRISPR screening, as well as earlier mapping efforts in *D. sechellia*/*D. simulans* hybrids (Amlou, et al. 1997; Amlou, et al. 1998; Jones 1998; Huang and Erezyilmaz 2015) revealed multiple genomic regions/genes contributing to the resistance to OA.

Considered separately, the experimental evolution and cell-based CRISPR screens identified many candidates. That there was very limited overlap between the screen results is unsurprising given their very different designs. The loss-of-function, cell-based screen of course has several limitations, as it cannot identify genes that are not expressed in the cell line, those essential for cell viability in the absence of OA, nor genes that function in biological processes only pertinent at a tissue/organismal level (e.g., cuticle biogenesis, neuronal signalling). Nevertheless, both lists are likely to contain multiple genes relevant to understanding OA susceptibility and resistance, and hence mode(s) of action of OA. Here, we focused on two resistance gene candidates, *kraken* and *Alkbh7*, which were common hits of both screens. We note, however, that neither of these genes were identified in previous studies of OA resistance (Amlou, et al. 1997; Amlou, et al. 1998; Jones 1998; Huang and Erezyilmaz 2015), which could reflect several factors, such as differences in OA toxicity assay conditions, developmental stages, species and molecular readouts (DNA or RNA), and the evident multilayered, polygenic nature of this trait (discussed further in (Marconcini, et al. 2026)). We note, however, that our recent GWAS of OA resistance in the founder panel of *D. simulans* used to establish the experimental evolution populations identified *bez* and CG13003 as candidate genes (Marconcini, et al. 2026), which showed significant CMH signals at generation 25 (but not generation 50) of the evolve-and-resequence experiment. (Neither gene could have been recovered in the CRISPR screen because they are not expressed in S2R+ cells (Hu, et al. 2017)).

The (partial) necessity, but insufficiency, of *kraken*, and lack of necessity but sufficiency of *Alkbh7* to confer OA resistance implies fundamentally different roles of these proteins. The expression of *kraken* in the gut and renal system supports a hypothesis of this protein as a detoxification enzyme, although future work will be necessary to determine whether Kraken has a direct role in OA degradation. The mild loss-of-function phenotype of *kraken* in both *D. melanogaster* and *D. sechellia* likely reflects substantial redundancy of defence mechanisms, including other potential detoxification mechanisms, especially given the plethora of enzymes expressed in the fly gut and Malpighian tubules (Lemaitre and Miguel-Aliaga 2013; Dow, et al. 2022). Indeed, extremely few associations between such enzymes and substrates and/or physiological roles have previously been defined. One exception is the Cytochrome P450 gene *Cyp6g1*, which has a key role in Malpighian tubules in conferring resistance to DDT (Yang, et al. 2007). Our data on Kraken therefore provide one of the first links between a natural toxic chemical and a putative detoxification enzyme.

The mitochondrial Alkbh7 enzyme likely has a distinct role in OA resistance. Insights from mammals are informative: mice lacking *Alkbh7* are obese, with higher body fat levels, proposed to result from a defect in metabolism of fatty acids (Solberg, et al. 2013). Molecularly, one study suggests that Alkbh7’s metabolic role stems from its regulation of RNA editing to control mitochondrial protein translation (Zhang, et al. 2021), although it is unclear whether this is its only function. While the *Alkbh7* loss-of-function phenotype is mild in drosophilids, it is striking that artificially elevated expression in *D. melanogaster* is sufficient to produce *D. sechellia*-like levels of OA resistance. This result implies a potent effect of Alkbh7 on the mitochondrial metabolic network to modulate toxin resistance, perhaps through control of fatty acid metabolism. While such ideas can be tested in future studies, we note that defects in lipid metabolism and mitochondrial function have been observed upon exposure to other types of insecticide (Martelli, et al. 2020; Martelli, et al. 2022). Intriguingly, our implication of Alkbh7 in the niche adaptation of *D. sechellia* has a potential parallel in bats, where this gene displays evidence of positive selection during dietary diversification of different species (Gutierrez-Guerrero, et al. 2020).

Taken together, our work has demonstrated the power of orthogonal selection approaches in the laboratory to understand toxin resistance, which might be relevant to understand a historic paradigm of this phenomenon (R’Kha, et al. 1991), as well as offering clues as to how the natural toxin OA exerts its lethal effects. Moreover, the methodology – notably cell-based CRISPR screening, which can now be extended to both gain-of-function approaches (Xia, et al. 2023) and mosquito cell lines (Viswanatha, et al. 2021) – is applicable to explore the molecular mechanisms of susceptibility and resistance of insects to the wide-range of artificial toxic chemicals used in the environment (Gandara, et al. 2024). Such knowledge could have application in the future characterization of novel types of insecticides.

## Author contributions

M.M. and R.B. conceived the project. M.M. designed and performed the experimental evolution experiment, generated and analysed all transgenic and mutant flies, and performed the comparative gene expression and molecular evolutionary analyses. S.C. generated the base populations and provided technical support for the experimental evolution experiment. M.M., R.B., R.V. and S.E.M. designed the cell screen. S.G. and R.V. tested and defined cell screen assay conditions. S.G. and M.B. performed the CRISPR/Cas9 screen and prepared DNA for sequencing. R.V. analysed the screen data, and M.M., R.B., R.V., S.E.M and N.P. contributed to interpretation of screen results. J.D. and C.R. performed histological experiments, supervised by D.H. M.M. and R.B. wrote the paper with input from other authors. All authors approved the final version of the manuscript.

## Acknowledgements

We thank Tadeusz Kawecki and Ana Marija Jakšić for advice on experimental evolution experiments, Tina Lenče and Jean-Yves Roignant for the gift of the *D. melanogaster Alkbh7* sgRNA transgenic line, the Bloomington *Drosophila* Stock Center (NIH P40OD018537) and the Vienna *Drosophila* Resource Center for *D. melanogaster* stocks, Christian Schlötterer and Neda Barghi for the *D. simulans* lines, Blaise Tissot-Dit-Sanfin for maintenance of *M. citrifolia* plants and Ambra Masuzzo for providing noni fruits. Figure icons were created with Biorender.com (https://BioRender.com/37yhdgj and https://BioRender.com/6j1dr4k). We are grateful to Tadeusz Kawecki, Philippe Reymond, Michael Shahandeh and members of the Benton laboratory for discussions throughout the project and comments on the manuscript. D.H. is funded by an ERC Starting Grant (101117267). N.P. is an Investigator of the Howard Hughes Medical Institute. Research in R.B.’s laboratory was supported by the University of Lausanne, an ERC Advanced Grant (833548) and the Swiss National Science Foundation (310030_219185 and 3200-0-239882).

## Data availability

Source data is provided in the supplementary data files. All other data, new materials and code are available from the corresponding author upon reasonable request.

## Materials and Methods

### *Drosophila* strains and husbandry

*Drosophila* stocks were maintained on standard wheat flour–yeast–fruit–juice medium under a 12 h light:12 h dark cycle at 25°C. Wild-type strains, as well as mutant and transgenic lines used in this study are listed in Table S1. Experimental evolution was performed using a subset of published *D. simulans* isogenic strains (Signor, et al. 2018) (Table S2).

### Generation of mutant and transgenic flies

#### Loss-of-function mutants

we generated *D. melanogaster* mutants for *kraken* and *Alkbh7* using in vivo CRISPR/Cas9 mutagenesis (Port, et al. 2020) by crossing flies expressing Cas9 in the germline (*nanos-Cas9*) to those expressing sgRNAs targeting the desired gene. For *kraken*, we used *P{hsFLP}1,y^1^,w^1118^;P{HD_CFD02326}attP40* as a source of sgRNAs driven by *Act5C-GAL4*; for *Alkbh7*, we used the CG14130 sgRNA line described in (Lenče 2021). Mutants were validated by genomic PCR and sequencing (Fig. S4B).

*D. sechellia* mutant flies were generated by WellGenetics Inc. using modified methods of (Kondo and Ueda 2013). For *kraken* (GM16772/LOC6617307) the upstream sequence CTGTAATGCTGGACGCGGAT[TGG] (PAM in square brackets) and the downstream sequence CCTGGTCACCCCGGATCGAG[TGG] were cloned separately into a U6 promoter plasmid. The 3xP3-RFP cassette, which contains a floxed 3xP3-RFP flanked by a 5’ homology arm (999 bp: from-1043 nt to-45 nt relative to the *kraken* ATG) and a 3’ homology arm (1063 bp: +1388 nt to +2450 nt) were cloned into *pUC57-Kan* as a donor template for repair. For *Alkbh7* (GM25300/LOC6605344) the upstream sequences ACTAATAAGAGTCGACCGGG[TGG], AACAGTCAAACAGCTGTTCC[TGG] and downstream sequences GTCCCAGTTTCAGGGTACGC[TGG], ATGGAAATGCGACGTGTCCT[GGG] were cloned separately into a U6 promoter plasmid. The 3xP3-RFP cassette, which contains a floxed 3xP3-RFP, 5’ homology arm (1037 bp:-1121 nt to-85 nt relative to the *Alkbh7* ATG) and 3’ homology arm (964 bp: +940 nt to +1903 nt) were cloned into *pUC57-Kan*. DNA plasmids encoding the sgRNAs and the corresponding homology-directed repair template were microinjected into embryos of *Dsec.07 pBAC(nos:Cas9,3P3-YFP)* (Auer, et al. 2020). In both cases, a single F1 line expressing the 3xP3-RFP selection marker was obtained, which was validated by genomic PCR and sequencing (Fig. S4B).

*D. melanogaster* and *D. sechellia* mutant flies were backcrossed for six generations to Canton-S and *Dsec*07, respectively.

*Transgenic flies*: to make the *UAS-Dmel\kraken* and *UAS-Dsec\kraken* transgenes, we amplified (using oligonucleotides listed in Table S6) the regions from the start to the stop codons from *w^1118^* and *Dsec07* genomic DNA and subcloned them into a *pUASTattB* vector (Bischof, et al. 2007). Transgenic flies were obtained by phiC31 integrase-mediated transgenesis into attP2 performed by BestGene Inc. and verified by sequencing.

### Toxin resistance assays

#### Tube assay

resistance to OA for the experimental evolution setup was scored adapting a previous protocol (Amlou, et al. 1997; Colson 2004). In brief, flies were lightly anesthetized with CO_2_ and ∼50 animals were placed in standard cultured tubes (25 mm diameter × 95 mm height, Milian SA) and provided with 1 ml of a 3% glucose solution on a tissue paper (Kimtech, 7552). 3 μl of pure OA (Sigma-Aldrich, CAS 124-07-2, C2875) were directly pipetted onto the tissue while the flies remained anesthetized. Direct contact with the acid was scrupulously avoided until all flies recovered. The fraction of alive flies was scored after 24 h. For the experimental evolution experiment, the surviving flies were collected (see below).

#### Plate assay

to measure OA resistance in the genetically-manipulated *D. melanogaster* and *D. sechellia* strains, we used a plate assay (Jones 1998). Compared to the experiments in tubes, this assay offers greater temporal resolution and higher dynamic range (in contrast to the experimental evolution experiment, we did not strive to maintain a certain fraction of surviving flies), as well as greater statistical power from tests of fewer animals. Ten female flies were lightly anesthetized with CO_2_ and placed in a Petri dish (60 mm diameter × 15 mm height) in which 2 μl of pure OA (or 30 μl for *D. sechellia*) were spotted in the centre of the underside of the lid. The flies were allowed to recover (typically <5 min), and immobile/dead were scored every 5 min for 1 h. The survival curves were calculated and plotted using the R package survminer (10.32614/CRAN.package.survminer). The statistical analysis was conducted fitting a mixed effect Cox regression model as implemented in the R package coxme (Therneau 2024). Genotype was included as a fixed effect, and replicate dishes were modelled as a random effect.

#### Noni resistance assay

noni fruit was collected from *M. citrifolia* plants grown in the University of Lausanne greenhouses. Five ripe (pale yellow) fruits were homogenized together, and the pulp puree frozen at-20°C. The noni assay was performed with the same setup as for tube assay described above, using 2 g of thawed noni puree placed on the bottom of the tube. Noni contains 3.06 g OA/kg fruit (Pino, et al. 2009); assuming an equivalent quantity in our fruits, 2 g noni pulp should contain 6.73 µl OA, approximately double the amount used for the experimental evolution between generations 1 to 25, and two thirds of the concentration during generations 30-50.

#### Long-term assay

To assess the long-term effects (i.e., beyond 24 h) of OA or noni pulp, two-day-old mated female flies were individually housed in vials containing either food supplemented with OA or 2g of noni puree. OA-supplemented food was prepared by thoroughly mixing OA into Formula 4-24® Instant Drosophila Medium, Blue (Carolina Biological Supply) to a final concentration of 0.9%. Flies were flipped into fresh food every two days, and mortality was monitored daily. Statistical analysis was run as for the plate assay.

#### Fecundity and developmental viability

10 mated adult female flies (4-7 days old) were allowed to lay eggs for 24 h on agar plates (2% agar, 2.5% sucrose in a 1:3 grape juice:water mix). The total number of eggs on each plate was counted. From each plate, 100 eggs were collected and transferred to standard fly food. Emerging adults were counted and removed daily until no further flies emerged.

#### Lifespan

Two-day-old mated female flies were individually housed in vials containing standard fly food and monitored daily until death. Fresh food was provided every 2–3 days.

#### Experimental evolution

*Selection procedure*: to generate the base populations for experimental evolution, we used 93 isogenic lines (Table S2) of the *D. simulans* panel established by (Signor, et al. 2018). We first crossed each of the lines together following a round-robin design to ensure equal contribution of each line to the starting genetic pool; 5 males and 5 females from each cross were then combined in ten independent replicates (Fig. S1D). These ten populations were maintained in bottles (57 mm diameter ×103 mm height, Milian SA) of standard fly food at high population size (∼1000 individuals per population) to minimize the effect of genetic drift. Each population was subdivided into smaller groups of ∼250 individuals per bottle (i.e., four bottles per population) to reduce competition. At each generation, individuals from the four bottles were mixed to prevent bottle-specific genetic drift. The experimental evolution experiment was initiated 10 generations after the creation of these populations.

From each population, 500 individuals (2-7 day old) were randomly collected and further split into 10 groups of 50 mixed-sex flies (i.e., ∼5000 flies total). Each group of 50 flies was then placed under selection with 3 μl of OA for 24 h (as described above); the survivors were collected and allowed to breed in food vials for 6-8 days, transferring them to fresh vials every 2-3 days. In the next generation, 500 flies per population were collected only from the last seeded vial, to maximize the probability of these flies being parented by the OA-resistant males and females and not offspring generated by females that mated before the selection procedure. These flies were then subjected to the same selection procedure. This process was repeated for 25 generations, and survivability was scored at each generation.

A second round of experimental evolution was performed by increasing the amount of OA to 9 μl between generations 31 and 50; this amount was determined empirically during generation 26-31 to re-establish the survivability close to the initial level (Fig. 1G). All successive generations were maintained following the same procedure.

As a control, the ten original populations were maintained in parallel in the same conditions as previously described (i.e. ∼1000 individuals per population subdivided into smaller groups of ∼250 individuals per bottle).

*DNA sequencing and analysis:* we pooled ∼200 individual flies per population per time point (generations 0, 25 and 50). Genomic DNA (and total RNA for qPCR, as described below) extraction was performed using the Quick DNA/RNA Miniprep plus kit (Zymo Research, D7003) following the manufacturer’s instructions. Samples were stored at-80°C.

We performed paired-end whole genome sequencing on an Illumina NovaSeq 6000 following Nextera DNA flex preparation, at an average coverage of >200×. Read quality was assessed with Fastqc (Wood 2011) and adapters were trimmed with Trimmomatic (Bolger, et al. 2014) providing the Nextera adapters sequences. We aligned the trimmed reads to the *D. simulans* reference genome (NCBI GCF_016746395.2) using bwa-mem (Li 2013) and the results were sorted and compressed using samtools (Danecek, et al. 2021). Duplicate reads were then removed with Picard MarkDuplicates (https://broadinstitute.github.io/picard/). Unplaced scaffolds and alignments with mapping quality <20 were removed with samtools. We then combined all the aligned sequences into a samtools pileup file and split this into individual chromosomes. Regions with zero coverage were removed with custom scripts. Sync files for each chromosome were generated with the mpileup2sync.pl script implemented in Popoolation2 (Kofler, et al. 2011) and used as input files for all the Evolve-and-Resequence analyses.

To explore the genetic variation of our samples we generated vcf files with vcftools 0.1.17 (Danecek, et al. 2011) and binary files with plink1.9 (Purcell, et al. 2007; Purcell 2020). We calculated the identity-by-state (IBS) matrix as implemented in plink1.9 (function-distance ‘ibs’).

*Evolve-and-Resequence analysis*: allele frequency changes between generation 0 and generations 25 or 50 were investigated via Popoolation2 and Bait-ER (Barata, et al. 2023).

We performed a Cochran–Mantel–Haenszel (CMH) test via the cmh-test.pl script implemented in Popoolation2 comparing base and evolved populations with parameters --min-count 12 and --min-coverage 50. We estimated a plausible significance threshold approximating the top 0.000005% of the simulated p-values based on the effect of genetic drift alone by running genome-wide forward simulations under neutrality in mimicree2 (Vlachos and Kofler 2018). We used the same genomic information from the original 93 lines as input and the average real survivability ratios observed throughout the selection experiment as –selection-regime parameter and 500 as –population-size parameter (Fig. S1B).

For Bait-ER, the input sync files were filtered to match --min-count 12 and --min-coverage 50 and scaled to a uniform coverage of 100 as in (Vlachos, et al. 2019) to avoid numerical problems due to high coverage. We ran the algorithm on each chromosome separately. Bait-ER models the effect of genetic drift directly using estimates of the effective population size (*N_e_*). We calculated *N_e_* estimates for each chromosome in each replicate population with the R package poolSeq (Taus, et al. 2017) and used the median value of each chromosome as “Population_size” parameter (Table S3). Other parameters included “Sampling correction” = 0 (i.e., binomial) and “Prior parameters” =0.001 (shape), 0.001 (rate). Data points were considered significant when the absolute natural logarithm of the Bayes Factor exceeded the conservative threshold of log(0.99/0.01) as in (Barata, et al. 2023).

The linkage disequilibrium decay in the 10 evolved populations was analysed across the whole genome using PopLDdecay (Zhang, et al. 2019). Linkage disequilibrium maps of focal regions were generated with LDBlockShow (Dong, et al. 2021) using the Solid Spine method based on D’ statistics originally introduced in Haploview (Barrett, et al. 2005).

For the genome-wide screen, all the genes within selection blocks were considered potential candidates. Based on the results on linkage disequilibrium decay, a block was arbitrarily defined as a genomic interval where at least two significant SNPs were found at a distance less than 50 kb. Consecutive SNPs respecting this definition were grouped into larger blocks and visualized as circus plots using the R package circlize (Gu, et al. 2014). The genes overlapping such blocks were extracted using BEDtools intersect (Quinlan and Hall 2010). Orthologous genes in *D. melanogaster* were retrieved using a combination of the OMA browser (Altenhoff, et al. 2021), DAVID (Huang da, et al. 2009; Sherman, et al. 2022), BLAST (Camacho, et al. 2009), NCBI (Sayers, et al. 2025) and FlyBase (Ozturk-Colak, et al. 2024), and subsequent manual curation.

### Genome-wide CRISPR/Cas9 screen in *D. melanogaster* S2R+ cells

*Library design, construction, and transfection*: design, synthesis and cloning of the genome-wide sgRNA library are previously described (Viswanatha, et al. 2025). To determine sub-lethal doses of OA appropriate for screening, the effect of a gradient of OA supplementation into media on attP+ Cas9+ cells was determined using CellTiter-Glo (Promega) following the manufacturer’s protocol; luminescence measurements were made using a Spectramax Paradigm (Molecular Devices).

For the screen, attP+ Cas9+ *D. melanogaster* S2R+ cells (RRID:CVCL_UD30) were transfected using Effectene and a 1:1 molar ratio of sgRNA library and pIntAC, as described (Viswanatha, et al. 2025). Briefly, transfected cells were distributed in 5 ml aliquots across ten 100 mm culture dishes and incubated overnight, then 5 ml of fresh media was added to each plate. After 4 days, cells were subjected to puromycin selection for 12 days, subculturing every 4 days, to establish a stable sgRNA-expressing cell library. For the selection assay, 2.5 × 10^7^ cells were plated onto four 15 cm diameter cell culture dishes to ensure sufficient sgRNA coverage, with each sgRNA represented approximately 1000 times (Viswanatha, et al. 2025).

*OA selection assay*: library-containing cells were left untreated or treated with either 0.075 μg/μl or 0.05 μg/μl OA (Sigma-Aldrich, CAS 124-07-2, C2875). Each replicate consisted of four 15 cm diameter cell culture dishes seeded with cells, achieving ∼1000× coverage. Two replicate sets were included for each of the two OA treatment conditions and one replicate for the untreated control. During the selection period, cells were split 1-2 times per week to avoid overgrowth. An aliquot of cells from each replicate was collected and stored for genomic prep following each split and at the end of the assay. The final samples were collected after 26 days of OA or control treatment.

*Genomic DNA extraction and sequencing*: genomic DNA was extracted from samples collected during the screen and at the final endpoint using a Zymo Quick-gDNA Miniprep kit (Zymo Research, D3025). DNA fragments containing the sgRNA sequences were amplified by PCR using what we calculated to be at least 1000 genomes per sgRNA as templates for each sample. PCR fragments were in-line barcoded using a previously described approach (Viswanatha, et al. 2018; Viswanatha, et al. 2019; Viswanatha, et al. 2025). Next-generation sequencing (Illumina NextSeq) was performed at the Biopolymers Facility at Harvard Medical School (RRID:SCR_007175). FASTQ sequence files were demultiplexed using in-line barcodes as sequence tags as described previously (Viswanatha, et al. 2018; Viswanatha, et al. 2025). All further processing was done using MAGeCK (v.0.5.4 and 0.5.9.4) (Li, et al. 2014) to generate readcount files and perform robust rank aggregation (RRA) analysis and Maximum likelihood estimation directly on the generated readcount files using default parameters. Plasmid readcounts were used to normalize RRA values.

After removing essential genes at a 2% FDR (2,517 genes; Data S4), we considered as candidates the top 5% genes overlapping across the two conditions that were over-represented or under-represented in the OA-exposed cell populations compared to the control cells (i.e., representing “susceptibility” and “resistance” genes, respectively). The gene-set enrichment analysis was performed with PANGEA 1.1 Beta using the *Drosophila* GO subsets (GO slim) and a p-value cut-off of 0.05 (Hu, et al. 2023).

### qPCR analysis

Reverse transcription of total RNA (extracted as described above) was performed with the PrimeScript first-strand cDNA synthesis kit (Takara, 6110A) following the manufacturer’s instructions. Samples were stored at-80°C.

Quantitative PCR experiments were conducted on a QuantStudio 6 Flex Real-Time PCR System using the PowerUp™ SYBR™ Green Master Mix following the manufacturer’s instructions. Each sample consisted of at least three biological replicates. We performed three technical replicates per reaction and removed replicates where the standard deviation was equal or greater than 0.5. We determined target specific amplification efficiencies for each gene (Table S4, Table S6). Relative quantification and statistical analysis were performed with the software qbase+ v3.4 (Hellemans, et al. 2007) using Calibrated Normalized Relative Quantities (CNRQs). Normalization was performed on the geometric mean of the relative quantities for the three most commonly-used reference targets: *Act5C*, α*Tub84B* and *RpL32* (Lu, et al. 2018). We assessed reference target stability by the geNorm expression stability value (M) and the coefficient of variation of the normalized reference gene relative quantities (CV) (Table S5) (Vandesompele, et al. 2002; Hellemans, et al. 2007).

### Population genetics analysis

Publicly available WGS data for 41 *D. sechellia* and 20 *D. simulans* genomes were downloaded from the Sequence Read Archive (PRJNA215932 and PRJNA395473) (Rogers, et al. 2014; Schrider, et al. 2018) with fasterq-dump (Leinonen, et al. 2010). FASTQ files were processed and aligned as described in the “Experimental evolution” section. Aligned files were phased with samtools phase (Danecek, et al. 2021) and the genes of interest were extracted with pilon (Walker, et al. 2014) to find variation among populations. CDS sequences were aligned based on amino acid information using translatorX (Abascal, et al. 2010). Gene sequences were aligned using Clustal Omega (Madeira, et al. 2024). McDonald-Kreitman test, Tajima’s D and Fu and Li’s statistics were calculated in DNAsp 6 (Rozas, et al. 2017). FUBAR analyses (Fast, Unconstrained Bayesian AppRoximation) were performed in HyPhy 2.5 (Kosakovsky Pond, et al. 2020). The consensus sequences of *D. simulans* and *D. sechellia* were computed by SnapGene (www.snapgene.com) “export consensus” with a >50% threshold. The sequence alignment and secondary structures were depicted using ESPript 3 (Robert and Gouet 2014).

### RNA fluorescence *in situ* hybridization

To detect *kraken* mRNA *in situ*, we used the RNAscope Multiplex Fluorescence Detection Reagents v2 (Advanced Cell Diagnostics, 323110) and RNAscope H_2_O_2_ and Protease Reagents (Advanced Cell Diagnostics, 322381). Surfaces and equipment were cleaned with RNAzap (Invitrogen, AM9780) prior to dissection. Intestinal tissues were dissected from *D. melanogaster* (Canton S) and *D. sechellia* (*D. sechellia* 07) females in RNAlater (Sigma-Aldrich, R0901) diluted 1:50 in DEPC-treated water and briefly rinsed in 1× PBS (prepared in ultrapure water) (ThermoFisher Scientific, 70011044). Dissected tissues were mounted on poly-L-lysine-coated slides in 1× PBS and fixed immediately in cold 4% formaldehyde (Sigma-Aldrich, F8775) for 30 min. The fixative was removed, and tissues were incubated with few drops of hydrogen peroxide for 10 min at room temperature (RT). Tissues were then washed 3× 5 min in wash buffer (Advanced Cell Diagnostics, 320058) preheated to 40°C. For tissue dehydration, slides were incubated with 200 μl of ethanol solutions in the following order: 50% ethanol for 5 min at RT, 70% ethanol for 5 min at RT and 100% ethanol for 5 min at RT (repeated twice). Slides were air-dried for 5 min, and samples incubated with few drops of Protease III for 30 min at RT.

To hybridise probes, tissues were washed in 1× PBS for 3× 5 min. 50 μl of C1 *(D. melanogaster-kraken-RA* (NM_057917), Advanced Cell Diagnostics, 1850601-C1); and 1 μl of C2 *(D. melanogaster-Rpl32* (NM_079843.4), Advance Cell Diagnostics, 533451-C2) RNAscope FISH probes were preheated to 40°C for 10 min, cooled at RT for 5 min, applied to tissues and the slides were placed in a humidified chamber for hybridization at 40°C for 2 h. Slides were hybridized with AMP reagents in sequential steps. A few drops of AMP1, warmed to RT, were applied to slides, which were incubated at 40°C for 30 min, followed by washes in wash buffer 3× 5 min. Next, a few drops of AMP2, warmed to RT, were applied to slides and incubated at 40°C for 30 min, followed by washes in wash buffer 3× 5 min. Finally, a few drops of AMP3, warmed to RT, were applied to slides, and incubated at 40°C for 15 min, followed by washes in wash buffer 3× 5 min.

A few drops of HRP-conjugated probe was applied for C1 and slides were incubated at 40°C for 15 min. After, rinsing in wash buffer 3× 5 min, Opal dyes 520 nm (Akoya Biosciences, OP-001001) and/or 570nm (Akoya Biosciences, OP-001003 (1:1000 in TSA multiplex buffer, Advanced Cell Diagnostics, 322809) were applied to slides and incubated at 40°C for 30 min. Slides were then washed and treated with HRP blocker (a few drops per slide) for 15 min at 40°C and washed in wash buffer for 3× 5 min. Tissues were counterstained with DAPI for 5 min at RT and washed once in the wash buffer, before mounting in Vectashield (Vector Laboratories, H1000).

A Leica STELLARIS-DIVE confocal microscope was used to generate all confocal images, using a water-immersive 40× objective or an oil-immersive 63× objective. The images were acquired using both HyDs as well as PMTs tailored for the fluorophores of each sample.

### Statistics and reproducibility

Data were analysed and plotted in R 4.0.3 (R Core Team 2021).

## Supplementary Information

### Supplementary Tables

**Table S1.** *Drosophila* strains.

| Genotype | Reference/Source |
| --- | --- |
| *D. sechellia* 07 (*Dsec07*) | *Drosophila* Species Stock Center (DSSC):14021-0248.07 |
| *D. simulans* 04 (*Dsim04*) | DSSC:14021-0251.004 |
| *D. melanogaster* Canton-S (*DmelCS*) |  |
| *D. sechellia* 28 (*Dsec28*) | DSSC: 14021-0248.28 |
| *D. simulans* 196 (*Dsim196*) | DSSC: 14021-0251.196 |
| *D. sechellia* Aride 3311 | (Shahandeh, et al. 2026) |
| *D. sechellia* Mahe 1311 | (Shahandeh, et al. 2026) |
| *D. sechellia* Praslin 7612 | (Shahandeh, et al. 2026) |
| *D. simulans* H1 | D. Matute |
| *D. simulans* LZV47 | D. Matute |
| *D. simulans* MD221 | D. Matute |
| *y*[1] *w*[*]; *P*{Act5C-GAL4-*w*}*E1*/CyO (*act-Gal4*) | RRID:BDSC_25374 |
| *y*[1] *w*[1118] (used as background for *act-Gal4*) | RRID:BDSC_6598 |
| *y*[1] *w*[67c23] (background of *Alkbh7<sup>o/e</sup>*) | RRID:BDSC_6599 |
| *y*[1] *w*[67c23]; *P*{*y*[+*mDint2*]<br>*w*[+*mC*]=*EPgy2*}*EY05855* (*Alkbh7<sup>o/e</sup>*) | RRID:BDSC_16679 |
| *y*,*w*[1118];*P*{attP,*y*[+],*w*[3']}(background of UAS-*kraken<sup>RNAi</sup>* and UAS-*Alkbh7<sup>RNAi</sup>*) | VDRC:v60100 |
| *P*{KK100356}*VIE-260B* (UAS-*kraken<sup>RNAi</sup>*) | VDRC:v105604 |
| *P*{KK102727}*VIE-260B* (UAS-*Alkbh7<sup>RNAi</sup>*) | VDRC:v109669 |
| *y*[1] *M*{*w*[+*mC*]=*nanos-Cas9.P*}*ZH-2A w*[*] (*nanos-cas9*) | RRID:BDSC_54591 |
| *P*{*hsFLP*}1, *y*1 *w*1118;<br>*P*{*HD_CFD02326*}attP40/CyO-GFP (*sgRNA* line for *Dmel**kraken* CRISPR mutant) | VDRC:v342579 |
| CG14130 *cris* (gRNA 1+2)/CyO (*sgRNA* line for *Dmel**Alkbh7* CRISPR mutant) | (Lenče 2021) |
| *y*[1] *w*[67c23];;UAS-*kraken* (*kraken<sup>o/e</sup>*) | This study |
| *y*[1] *w*[67c23];;UAS-*Dsec**kraken* (*Dsec**kraken<sup>o/e</sup>*) | This study |
| *y*[1] *w*[67c23]; *P*{*y*[+*t7.7*]=*CaryP*}attP2 (background of *kraken<sup>o/e</sup>*) | This study |
| *Dmel**kraken*<sup>1</sup> | This study |
| *Dmel**Alkbh7*<sup>1</sup> | This study |
| *Dsec**kraken*<sup>RFP</sup> | This study |
| *Dsec**Alkbh7*<sup>RFP</sup> | This study |

**Table S2.** *D. simulans* lines used to generate the base populations. *-see separate Excel file*.

**Table S3.** Effective population size estimates from the experimentally-evolved populations.

|  | 2L | 2R | 3L | 3R | 4 | X |
| --- | --- | --- | --- | --- | --- | --- |
| pop1 | 80.65681 | 101.5796 | 107.7145 | 100.5748 | 369.3642 | 107.3616 |
| pop2 | 135.7243 | 56.29977 | 76.40593 | 91.41878 | 333.638 | 53.66758 |
| pop3 | 74.08474 | 119.4491 | 95.20849 | 128.4216 | 193.5982 | 99.40552 |
| pop4 | 83.05221 | 89.62312 | 112.3929 | 116.7979 | 171.2875 | 112.1581 |
| pop5 | 123.0386 | 106.8179 | 107.448 | 81.31624 | 364.4287 | 83.55315 |
| pop6 | 98.8018 | 77.03804 | 60.71772 | 147.1788 | 124.9405 | 75.48775 |
| pop7 | 119.4038 | 101.4515 | 96.22689 | 114.8849 | 431.7489 | 114.6792 |
| pop8 | 136.6513 | 176.4308 | 70.8519 | 130.9822 | 639.7962 | 78.70108 |
| pop9 | 95.58733 | 101.7351 | 122.8352 | 78.80166 | 109.3757 | 69.50865 |
| pop10 | 141.862 | 101.6789 | 107.512 | 83.0109 | 376.8733 | 71.34583 |
| mean | 108.8863 | 103.2104 | 95.73134 | 107.3388 | 311.5051 | 86.58684 |
| median | 109.1028 | 101.6293 | 101.8374 | 107.7299 | 349.0333 | 81.12711 |

**Table S4.** qPCR primer efficiency.

| Gene | Efficiency | SE |
| --- | --- | --- |
| *kraken* | 1.998 | 0.041 |
| *Alkbh7* | 1.977 | 0.016 |
| *Act5C* | 1.976 | 0.022 |
| *αTub84B* | 2.027 | 0.023 |
| *RpL32* | 2.053 | 0.031 |

**Table S5.** geNorm analysis of reference target stability. *-see separate Excel file*.

**Table S6.** Oligonucleotides. *-see separate Excel file*.

## Supplementary Figure legends

**Fig. S1.**
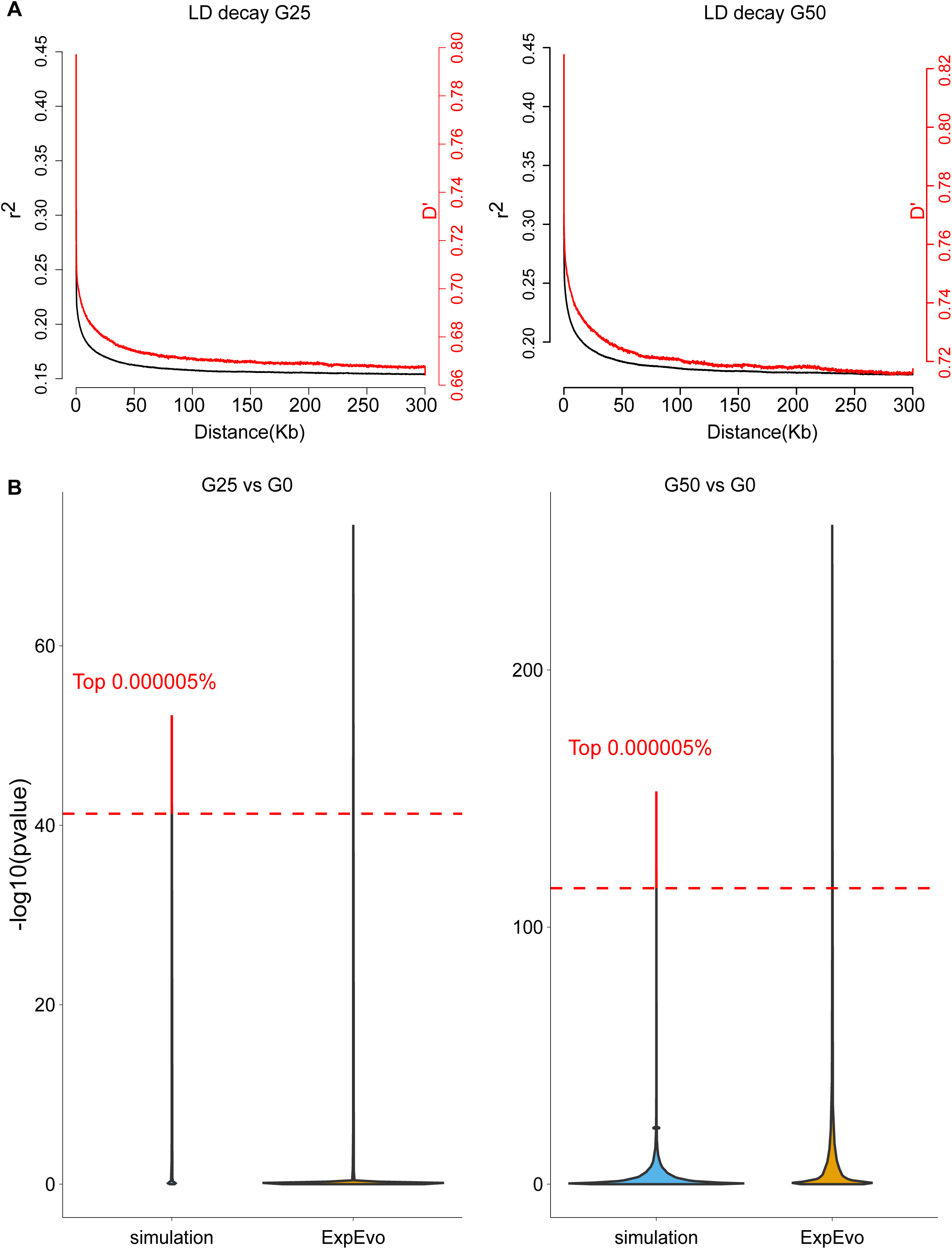
Linkage disequilibrium decay, and comparison of simulated and observed p-value distributions in the experimentally evolved populations. **A** Linkage disequilibrium decay at generations 25 (left) and 50 (right). Black curves represent the trajectory of r2 values (y-axis left) as genomic distances increase (x-axis). Red curves represent the same but for D’ values (y-axis right). **B** Violin plots showing the distribution of p-values from the CMH test, comparing allele frequency changes between generation 25 and 0 (left), and generation 50 and 0 (right). Blue violins represent p-values derived from forward evolution simulations, while orange violins correspond to p-values from experimentally evolved populations. Red dashed lines mark the top 0.000005% of simulated p-values, serving as the significance threshold for the experimental data.

**Fig. S2.**
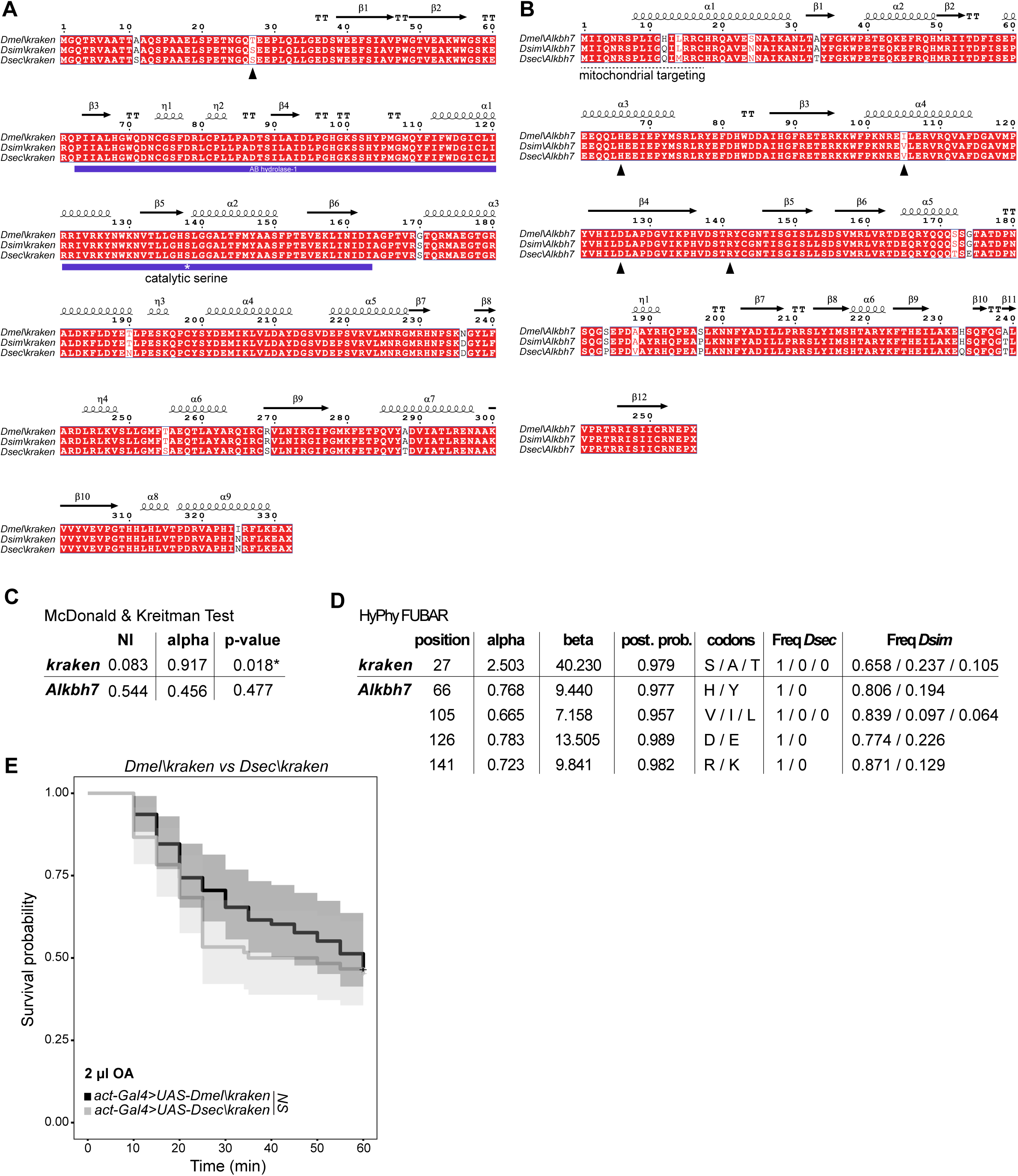
Sequence and molecular evolutionary analyses of *kraken* and *Alkbh7*. A,B. Protein sequence alignment and secondary structure of Kraken (**A**) and Alkbh7 (**B**) from the reference sequence of *D. melanogaster* (NP_477265.1 and NP_648511.2) and the consensus sequences of *D. simulans* and *D. sechellia*. Sequence data from PRJNA215932 and PRJNA395473 (see Materials and Methods). Black triangles indicate the residues where signs of selection were detected by the FUBAR analysis (see **D**). **C** Results from the McDonald-Kreitman test. NI = neutrality index; alpha = proportion of substitutions driven by positive selection; p-value = p-value resulting from a two-tailed Fisher’s exact test. **D** FUBAR analysis to detect pervasive selection as implemented in HyPhy. Position = codon position; alpha = mean posterior synonymous substitution rate; beta = mean posterior nonsynonymous substitution rate; posterior prob = posterior probability of positive selection; codons = alternative codons, Freq Dsec = frequency of the alternative codons in *D. sechellia* in the same order as codons; Freq Dsim = frequency of the alternative codons in *D. simulans* in the same order as codons. **E** Survival curves of *D. melanogaster* overexpressing *Dmel\kraken* and *Dsec\kraken* exposed to 2 μl OA. Each replicate consisted of 10 2-7-day-old female flies. Genotypes: *act-Gal4/+;UAS-Dmel\kraken/+*, *act-Gal4/+;UAS-Dsec\kraken/+*. N replicates ≥6, n flies ≥60. Shading indicates 95% confidence intervals. Significance is based on the resulting mixed effect Cox regression model. NS p>0.05. Raw data in Data S5.

**Fig. S3.**
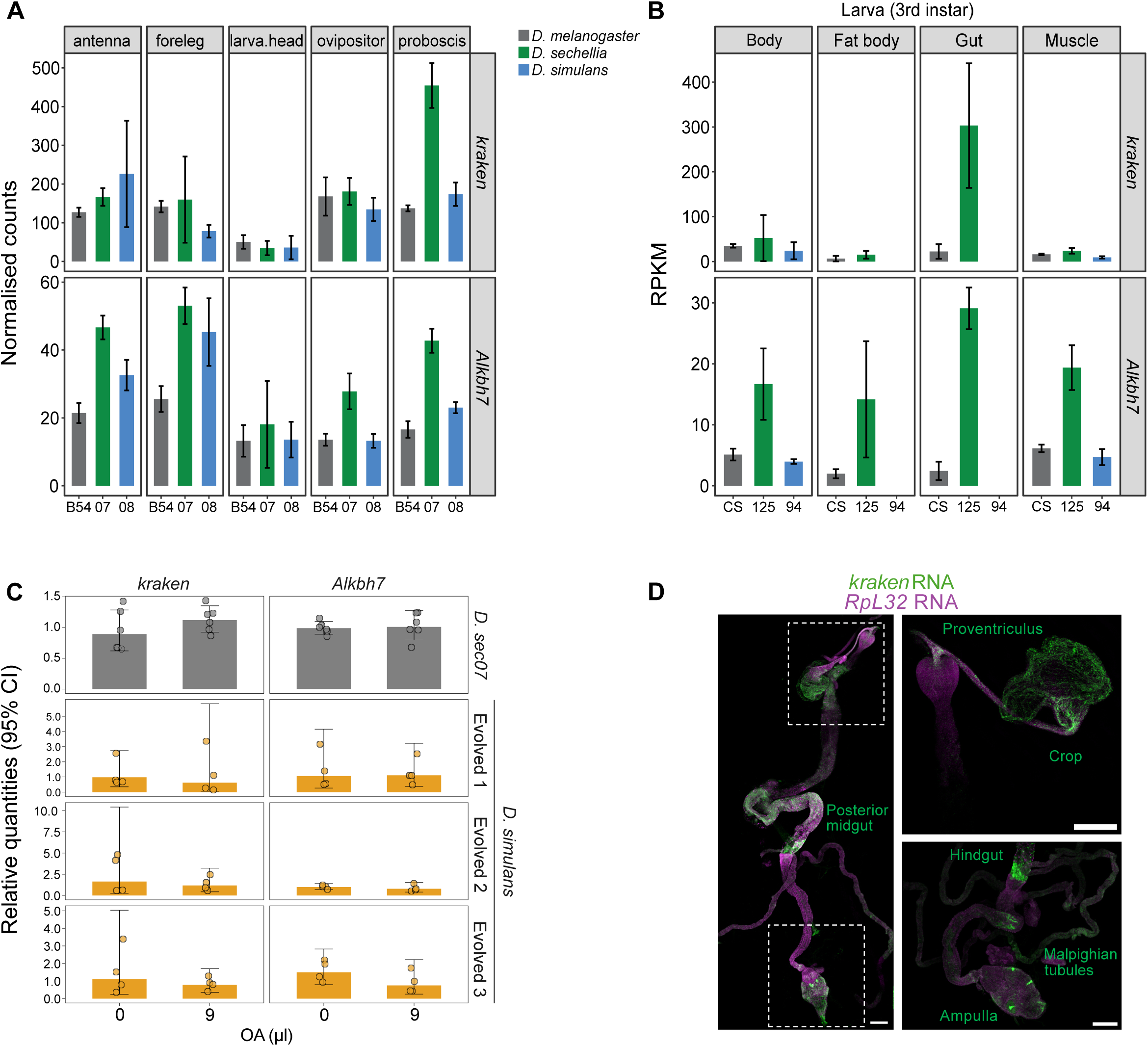
Additional expression analyses of *kraken* and *Alkbh7*. **A** Bar plots showing the expression levels of *kraken* and *Alkbh7* across various adult *Drosophila* tissues and species. Data from (Bontonou, et al. 2024). **B** Bar plots showing the expression levels of *kraken* and *Alkbh7* across various larval *Drosophila* tissues and species. Data from (Watanabe, et al. 2019) (“medium diet” condition). **C** Bar plots showing the effect of exposure to 9 μl OA (in the tube assay) on *kraken* and *Alkbh7* expression in *D. sechellia* and three evolved (G50) *D. simulans* populations. **D** Left: RNAscope detection of *kraken* (green) and *Rpl32* (magenta) transcripts in the gut and Malpighian tubules. Right: higher-magnification showing *kraken* transcript expression in the indicated tissues. Scale bars, 200 μm. This expression pattern was observed in tissues from >10 individuals.

**Fig. S4.**
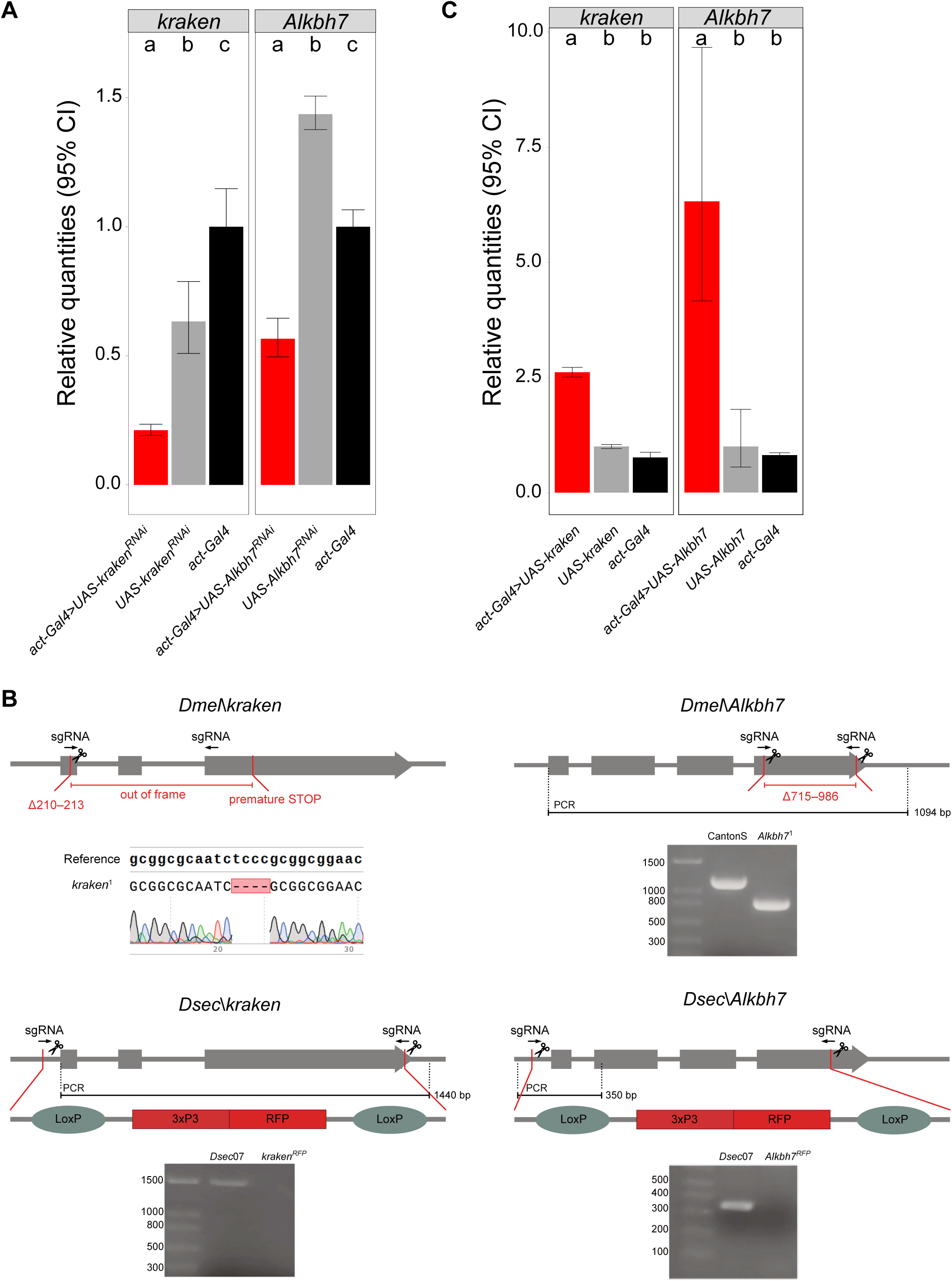
Validation of RNAi and overexpression lines and generation of mutants for *kraken* and *Alkbh7*. **A** Quantitative RT-PCR analysis of RNAi-mediated knockdown of *kraken* and *Alkbh7* in *D. melanogaster*. Genotypes as in Fig. 5A. **B** Schematics of the CRISPR/Cas9 design to generate *D. melanogaster* and *D. sechellia* mutants for *kraken* and *Alkbh7*. Validation was performed by PCR and Sanger sequencing. The *Dse*c\*kraken^RFP^* mutant comprised a 1431 bp deletion (-44 to +1,387 nt relative to the ATG), replaced with the 3xP3-RFP cassette. The *Dse*c\*Alkbh7^RFP^* mutant was a 1023 bp deletion (-84 nt to +939 nt relative to the ATG), replaced with the 3xP3-RFP cassette. **C** Quantitative RT-PCR analysis of Gal4/UAS-mediated overexpression of *D. melanogaster kraken* and *Alkbh7*. Genotypes as in Fig. 5C.

**Fig. S5.**
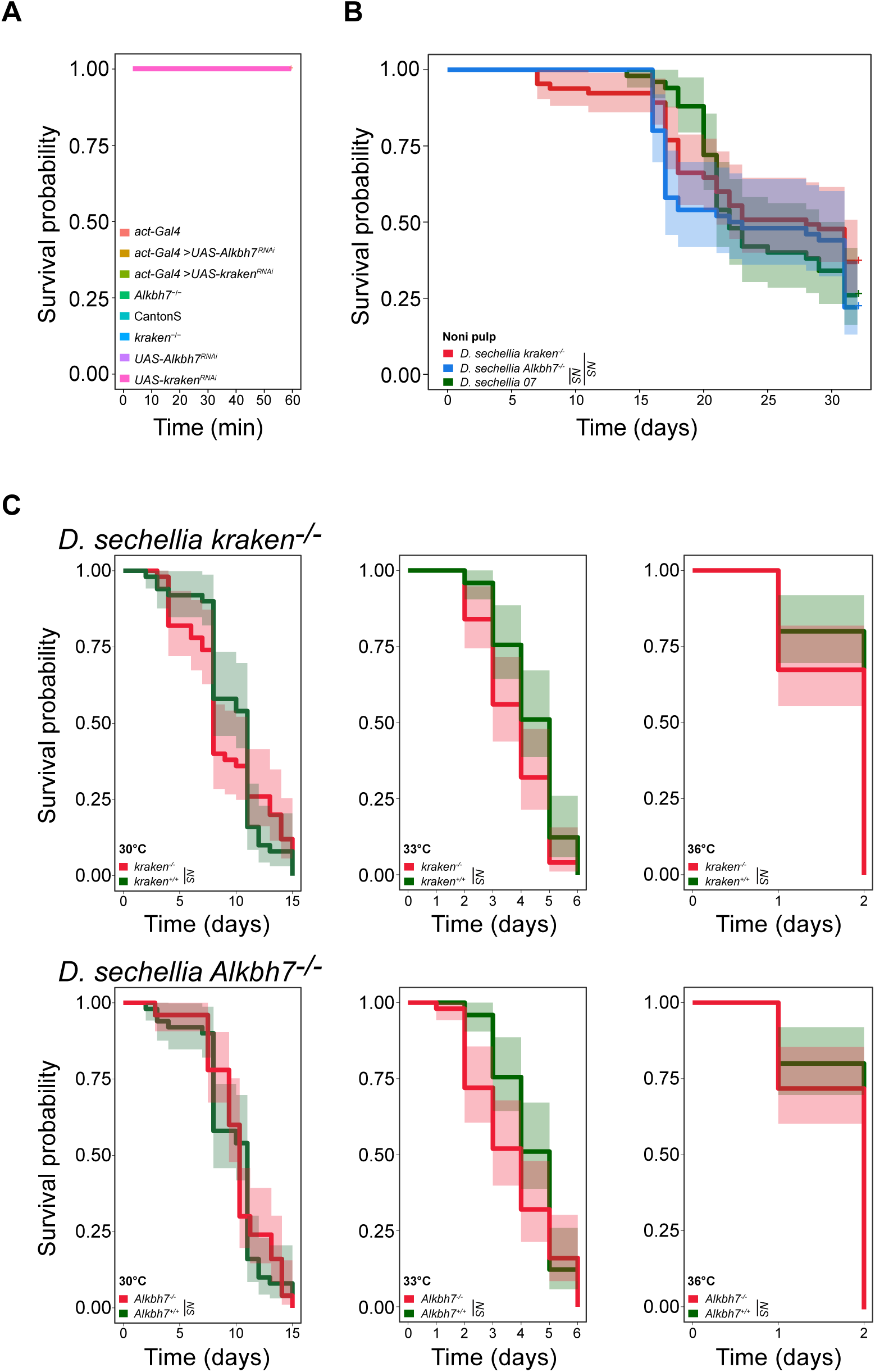
Control survival curves in the absence of OA. **A** Survival curves of adult flies of the indicated genotypes tested in the plate assay without OA. Each replicate consisted of 10 2-7-day-old female flies. N replicates ≥5, n flies ≥50. All genotypes display full viability, rendering datapoints invisible beneath the topmost displayed genotype (*UAS-kraken^RNAi^*). Genotypes as in Fig. 5A-B. **B** Survival curves of wild-type (*Dsec07*), and *kraken* (*Dsec\kraken^RFP^*) and *Alkbh7* (*Dsec\Alkbh7^RFP^*) mutant *D. sechellia* on homogenized noni pulp. N = 50 per strain. Significance is based on the resulting mixed effect Cox regression model. NS p>0.05. **C** Survival curves of *kraken* (*Dsec\kraken^RFP^*, top) and *Alkbh7* (*Dsec\Alkbh7^RFP^*, bottom) mutant *D. sechellia* compared to wild-type (*Dsec07*) flies at 30, 33, 36 degrees Celsius. N ≥ 50 per strain/condition. Statistics as in **B**.

**Data S1.** Source data for Fig. 1

**Data S2.** Experimental evolution data available at: https://drive.google.com/file/d/1Gy3k9eYvQ8xQMjIsTCqWevRaO_b1_JY/view?usp=sharing

**Data S3.** Source data for Fig. 2

**Data S4.** Source data for Fig. 3

**Data S5.** Source data for Fig. S2

**Data S6.** Source data for Fig. 4

**Data S7.** Source data for Fig. 5 and Fig. S5.

